# Oxidative phosphorylation is required for powering motility and development of the sleeping sickness parasite *Trypanosoma brucei* within the tsetse fly vector

**DOI:** 10.1101/2021.06.30.450569

**Authors:** Caroline E. Dewar, Aitor Casas-Sanchez, Constentin Dieme, Aline Crouzols, Lee R. Haines, Álvaro Acosta-Serrano, Brice Rotureau, Achim Schnaufer

**Affiliations:** Institute of Immunology and Infection Research, University of Edinburgh, United Kingdom; Department of Vector Biology, Liverpool School of Tropical Medicine, United Kingdom; Trypanosome Transmission Group, Trypanosome Cell Biology Unit, Institut Pasteur and INSERM U1201, Paris, France; Department of Tropical Disease Biology, Liverpool School of Tropical Medicine, United Kingdom

**Author notes:** Department of Chemistry, Biochemistry and Pharmaceutical Sciences, University of Bern, Bern, Switzerland (Caroline E. Dewar). Corresponding authors: (AAS), (BR), (AS). these authors contributed equally to this work (Aitor Casas-Sanchez, Constentin Dieme). these authors also contributed equally to this work (Álvaro Acosta-Serrano, Brice Rotureau, Achim Schnaufer).

## Abstract

The single-celled parasite *Trypanosoma brucei* causes sleeping sickness in humans and nagana in livestock and is transmitted by hematophagous tsetse flies. Lifecycle progression from mammalian bloodstream form to tsetse midgut form and, subsequently, infective salivary gland form depends on complex developmental steps and migration within different fly tissues. As the parasite colonises the glucose-poor insect midgut, its ATP production is thought to depend on activation of mitochondrial amino acid catabolism via oxidative phosphorylation. This process involves respiratory chain complexes and the F_1_F_O_-ATP synthase, and it requires protein subunits of these complexes that are encoded in the parasite’s mitochondrial DNA (kinetoplast or kDNA). Here we show that a progressive loss of kDNA-encoded functions correlates with an increasingly impaired ability of *T. brucei* to initiate and complete its development in the tsetse. First, parasites with a mutated F_1_F_O_-ATP synthase with a reduced capacity for oxidative phosphorylation can initiate differentiation from bloodstream to insect form, but they are unable to proliferate *in vitro*. Unexpectedly, these cells can still colonise the tsetse midgut. However, these parasites exhibit a motility defect and are severely impaired in colonising or migrating to subsequent tsetse tissues. Second, parasites with a fully disrupted F_1_F_O_-ATP synthase complex that is completely unable to produce ATP by oxidative phosphorylation can still differentiate to the first insect stage *in vitro* but die within a few days and cannot establish a midgut infection *in vivo*. Third, mutant parasites lacking kDNA entirely can initiate differentiation but die within 24 h. Together, these three scenarios show that efficient ATP production via oxidative phosphorylation is not essential for initial colonisation of the tsetse vector, but it is required to power trypanosome migration within the fly.

## Introduction

The protist parasite *Trypanosoma brucei* is the causative agent of human African trypanosomiasis, also called sleeping sickness [1]. The life cycle of *T. brucei* is complicated in that it involves at least nine distinct stages within the mammalian bloodstream and tissues and the tsetse fly vector [2–4]. The distinct biology of these life cycle stages is closely linked to changes in energy metabolism and regulation of the activity of respiratory chain and F_1_F_O_-ATP synthase complexes in the inner membrane of the parasite’s single mitochondrion. As in other eukaryotes, these complexes are comprised of both nuclearly and mitochondrially encoded subunits, and regulation of their expression therefore requires tight coordination between these organelles. In kinetoplastids such as *T. brucei*, structure and expression of the mitochondrial genome, termed kinetoplast DNA or kDNA, is extraordinarily complex. *T. brucei* kDNA comprises dozens of maxicircles - the equivalent of mitochondrial DNA in other eukaryotes - and thousands of minicircles [5]. The latter encode guide RNAs (gRNAs) for remodelling of 11 of the 18 maxicircle-encoded mRNAs by uridylyl insertion and deletion, a maturation process called RNA editing [6].

In the proliferative long slender bloodstream form (BSF), ATP production is non-mitochondrial via glycolysis, exploiting the glucose-rich environment within the mammalian bloodstream [7, 8]. The BSF mitochondrion lacks cyanide-sensitive oxygen uptake: respiratory complexes III (ubiquinol:cytochrome c oxidoreductase) and IV (cytochrome oxidase) are absent, and the corresponding maxicircle- and nuclearly-encoded transcripts are only present at very low levels [9–13]. The BSF mitochondrion also lacks or has less developed cristae, which are invaginations of the inner mitochondrial membrane that are ultrastructural hallmarks of oxidative phosphorylation [14, 15]. Despite the absence of cytochrome-mediated respiration, BSF parasites are dependent on oxygen for survival since an alternative oxidase (AOX) in the inner mitochondrial membrane is critically involved in maintaining the redox balance for glycolysis [16]. BSF cells use the F_1_F_O_-ATP synthase complex in reverse: ATP hydrolysis by the F_1_ part provides the energy for pumping protons from the mitochondrial matrix through the F_o_ part into the intermembrane space. This activity generates a mitochondrial membrane potential, ΔΨm, essential for protein import and exchange of metabolites and therefore fundamental for mitochondrial function [17–19].

Long slender BSF *T. brucei* differentiate to the cell cycle-arrested stumpy BSF upon reaching high parasite numbers, induced by quorum sensing [20–22]. The stumpy form is pre-adapted to survival in the tsetse fly vector by being more resistant to pH and proteolytic stresses, and by beginning to prepare its mitochondrial energy metabolism for the change in nutrient availability. This adaptation includes upregulation of mitochondrial ATP production capability via substrate level phosphorylation and beginning of cristae formation [20,23–25].

Once ingested with a bloodmeal, stumpy forms differentiate into proliferative procyclic forms (PCF) within the fly midgut. Mitochondrial ATP production is vital to the PCF [26–28]. Its inner mitochondrial membrane is now invaginated to form prominent cristae and contains complexes I (NADH:ubiquinone oxidoreductase), II (succinate dehydrogenase), III and IV of a classical cytochrome-mediated respiratory chain, along with complex V (F_1_F_O_-ATP synthase) [29, 30]. In the tsetse fly midgut, proline and intermediates of the tricarboxylic acid cycle are thought to be the main energy sources [31–35], with glucose only transiently abundant after bloodmeals [36]. In glucose-depleted *in vitro* growth conditions, which are similar to the tsetse fly midgut environment, ATP production via oxidative phosphorylation is essential [28, 37]. In glucose-rich *in vitro* growth conditions, only ATP production by mitochondrial substrate level phosphorylation appears to be essential in PCF [26, 27]; however, the electron transport chain remains essential under these conditions [27, 38]. Complexes III and IV, despite being structurally divergent [9,39–43], are thought to have classical enzymatic roles in the PCF respiratory chain: by pumping protons from the mitochondrial matrix into the intermembrane space they generate a proton motive force and a ΔΨm upon the uptake of oxygen [44, 45]. Under glucose-rich conditions, it is possibly this generation of ΔΨm that becomes the sole essential function of complexes III and IV. Regardless of whether production of mitochondrial ATP involves oxidative or substrate level phosphorylation, it requires the ADP/ATP carrier (AAC) to function in the direction of ATP export from the mitochondria to fuel processes in other parts of the cell [46].

By around day 3-7 post infection, PCF are found in the ectoperitrophic space of the midgut, which, according to a long-standing model, they have reached by crossing the peritrophic matrix (PM) in the anterior midgut [4]. However, recent findings suggest that *T. brucei* colonises the PV’s ectoperitrophic space by crossing the immature, fluid-like PM before it hardens [47]. Within this revisited model, procyclic trypanosomes may colonise first the anterior midgut or establish parallel infections at both the anterior midgut and PV [47–51]. Regardless of the topology of the infection, approximately after two weeks parasites differentiate into long epimastigotes within the PV lumen, a process that involves the reversal of the positions of the kinetoplast and nucleus [52]. Long epimastigotes then divide asymmetrically into long and short epimastigotes, aiding the delivery of short, less motile epimastigotes to the salivary glands [53, 54]. Few parasites are able to complete this complex migration [55], and only a small proportion (∼10%) of experimentally infected tsetse flies are found with infected salivary glands [56]; in the field, salivary gland infection rates are even lower compared to experimentally infected ones (< 1%) [57–59]. Once in the salivary glands, epimastigotes attach to the epithelium, proliferate and thus colonise these tissues [4, 60]. Epimastigotes undergo further asymmetric divisions to continuously yield trypomastigotes that are able to develop into cell cycle-arrested metacyclics around 3-4 weeks post infection [61]. After injection into the mammalian host upon a tsetse bite, metacyclics develop into BSF to begin the life cycle again. Much less is known about potential changes in mitochondrial energy metabolism after the PCF parasites have left the midgut, owing to the difficulty of culturing other insect stages in the laboratory. Recently, it was discovered that overexpression of the regulatory RNA binding protein RBP6 in PCF triggers efficient differentiation into metacyclic forms *in vitro* [62]. Using this experimental system, a switch from cytochrome-based to AOX-mediated respiration and a general retrogression of mitochondrial structure in metacyclic forms were demonstrated [62, 63].

The kDNA encodes essential components of the multi-subunit complexes involved in oxidative phosphorylation or in the translation of these components: eight subunits of complex I (ND1-5, 7-9), subunit apocytochrome b of complex III, three subunits of complex IV (COXI-III), subunit A6 of complex V, two hydrophobic protein subunits of the mitoribosome, S12 (RPS12) and S3 (MURF5), and the two mitoribosomal rRNAs [64, 65]. As discussed above, complex V is essential in both BSF and PCF, and complexes III and IV are essential in PCF. As a consequence, mitochondrial gene expression is critical for survival of both BSF and PCF *T. brucei*, although with PCF this has only been formally assessed *in vitro* [66–69]. (The role of complex I in energy metabolism in either the BSF or PCF is uncertain; this has been debated extensively elsewhere [24,70–77]).

Intriguingly, ‘akinetoplastic’ or ‘dyskinetoplastic’ BSF *T. brucei* cells that lack all (kDNA^0^) or critical parts of kDNA (kDNA^-^), respectively, occur in nature as subspecies *T. b. evansi* and *T. b. equiperdum* and have also been generated in the lab [64,78–80]. In nature, these kDNA mutants are directly transmitted from mammal to mammal, either by mechanical transfer via successive fly bites or sexually, i.e., without any cyclical development or proliferation in an insect vector. This apparent contradiction regarding the essentiality of kDNA in BSF *T. brucei* was resolved by the discovery of point mutations in the nuclear-encoded F_1_-ATPase subunit γ in akinetoplastic and dyskinetoplastic forms, for example a change of leucine in position 262 to proline (L262P). When introduced into wild type BSF, these mutations can compensate for kDNA loss by enabling cells to manufacture ΔΨm independently of the F_o_ part of the ATP synthase [17,66,81]. Therefore, the F_o_ subunit A6 is no longer essential (and neither are, consequently, the mitoribosomal protein and rRNA subunits), and the cells can survive as BSF without any kDNA gene products.

The molecular mechanism of the compensatory mutations in subunit γ is not fully understood, but they enable F_O_-independent generation of ΔΨm via the electrogenic action of AAC-mediated ATP^4-^/ADP^3-^ exchange across the inner mitochondrial membrane [17,24,66,82]. Kinetic studies with yeast expressing mutated γ subunits have demonstrated a lowered K_m_ value for ATP in the F_1_-ATPase reaction, suggesting that the mutant complex has a higher affinity for its substrate [17, 83]. More efficient F_1_-mediated ATP hydrolysis, perhaps in the vicinity of the AAC, might support this essential electrogenic ATP^4-^/ADP^3-^ exchange [17, 83].

On the other hand, the mutations appear to impair the function of the F_1_F_O_ enzyme when it is required to operate in the direction of ATP synthesis. Physiological and structural studies in yeasts have indicated that similar mutations in the yeast ATP synthase γ subunit alter its interactions with the catalytic F_1_-β subunit by partially uncoupling the F_1_ moiety from the F_o_ moiety [84–89]. In the presence of an active electron transport chain, impaired coupling of F_1_ and F_O_ action will result in some dissipation of ΔΨm and, consequently, diminished efficiency of the ATP synthesis reaction.

In this study we investigated the consequences of compensatory F_1_-γ mutations and kDNA loss for life cycle progression of *T. brucei* in the tsetse fly vector. Available evidence points to kDNA being essential for PCF viability and fly transmissibility [90]. RNAi knockdown of genes required for kDNA replication and division was lethal in PCF *in vitro* [69,91–93], and chemically induced kDNA^0^ PCF cells were also non-viable *in vitro* [68, 94]. Similarly, from a mixed population of kDNA^+^ and kDNA^0^ parasites (the latter were obtained by treatment of kDNA^+^ parasites with acriflavine), only kDNA^+^ cells were able to establish a tsetse midgut infection [95]. Finally, *T. b. evansi* and *T. b. equiperdum* isolates have been documented as being unable to transform into PCF *in vitro* [81,96,97]. A caveat with these studies was that they had been performed either with dying parasites unable to survive as BSF, or with ‘monomorphic’ strains that were unable to develop into the differentiation competent stumpy forms. The ability of viable, ‘pleomorphic’ kDNA^0^ strains to infect and potentially develop in the tsetse fly vector has never been formally assessed.

Here, we have built on our development of a range of pleomorphic BSF mutants that represent various defects in the expression of kDNA-encoded functions (Table 1) to assess the role of kDNA in stumpy to PCF differentiation, in the proliferation of PCF *in vitro*, and in the completion of the parasitic life cycle in tsetse flies.

**Table 1.**
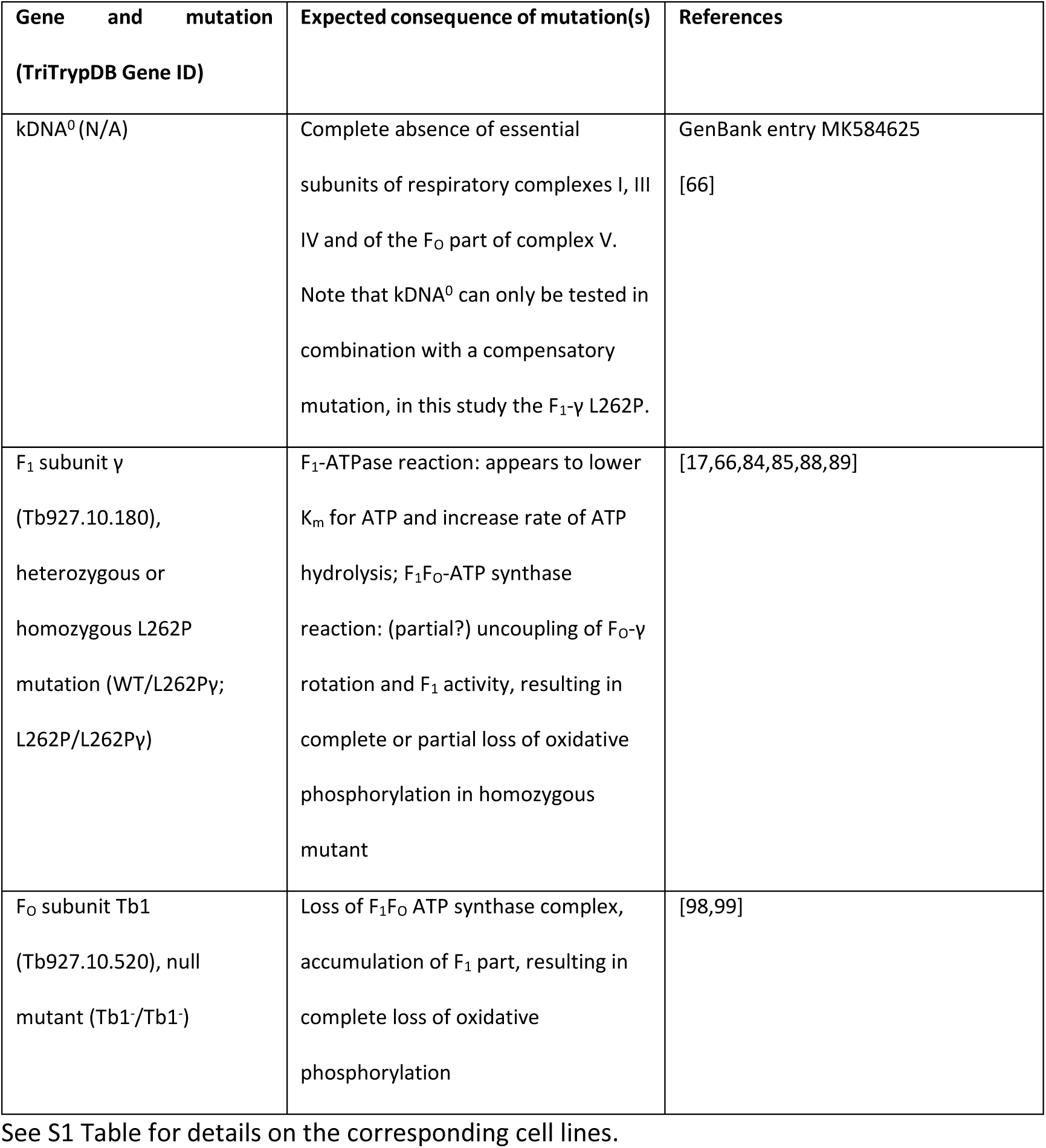
Mutations affecting trypanosome mitochondrial functions that were assessed for their effects on tsetse infectivity in this study.

## Results

To investigate the requirement for functional kDNA and a fully functional F_1_F_O_-ATP synthase in the differentiation of *T. brucei* from stumpy BSF to PCF we generated a set of mutant cell lines (Table 1 and S1 Table). In the pleomorphic cell line EATRO 1125 (AnTat1.1 90:13) we first replaced one allele or both alleles of the nuclear-encoded F_1_F_o_-ATP synthase subunit γ with either a wild type version (WTγ; serving as control), or a version with the L262P mutation (L262Pγ), as detailed in [24]. This resulted in cell lines WT/WTγ, WT/L262Pγ and L262P/L262Pγ, respectively (Table 1 and S1 Table). The L262Pγ mutation, in homozygous or heterozygous configuration, enables slender BSF *T. brucei* to survive and proliferate without kDNA [66]. We generated two cell lines lacking kDNA (kDNA^0^ #1 and #2) by treating two distinct clones of genotype WT/L262Pγ (#1 and #2) with the kDNA intercalating dye acriflavine [24]. The absence of kDNA in these two cell lines, and the lack of an *in vitro* growth phenotype in absence of kDNA, have been shown previously [24]. Moreover, *T. brucei* kDNA^0^ parasites can still differentiate to the stumpy BSF [24].

To test the requirement of kDNA for differentiation to PCF, we harvested stumpy kDNA^0^ and control cells from infected mice, and induced differentiation into PCF *in vitro* by adding 6 mM *cis*-aconitate (CA) to the medium and shifting the temperature from 37°C to 27°C for 24 h [100–104]. By microscopic inspection, no intact and motile kDNA^0^ cells were visible at this point, with only non-motile cells and debris remaining (data not shown). This was in contrast to all other cell lines which remained motile, although some debris was present. We then harvested cultures by centrifugation and assessed lysates by western blotting for expression of EP procyclin, an early marker for PCF differentiation [105–107] (Fig 1A). Compared to WT/WTγ and WT/L262Pγ lysates, the EP procyclin signal was much reduced in material recovered from kDNA^0^ cultures. These results suggest that kDNA^0^ BSF cells differentiated to PCF but died soon afterwards, which implies that the requirement for kDNA in PCF cells cannot be compensated for by the L262Pγ mutation.

**Figure 1.**
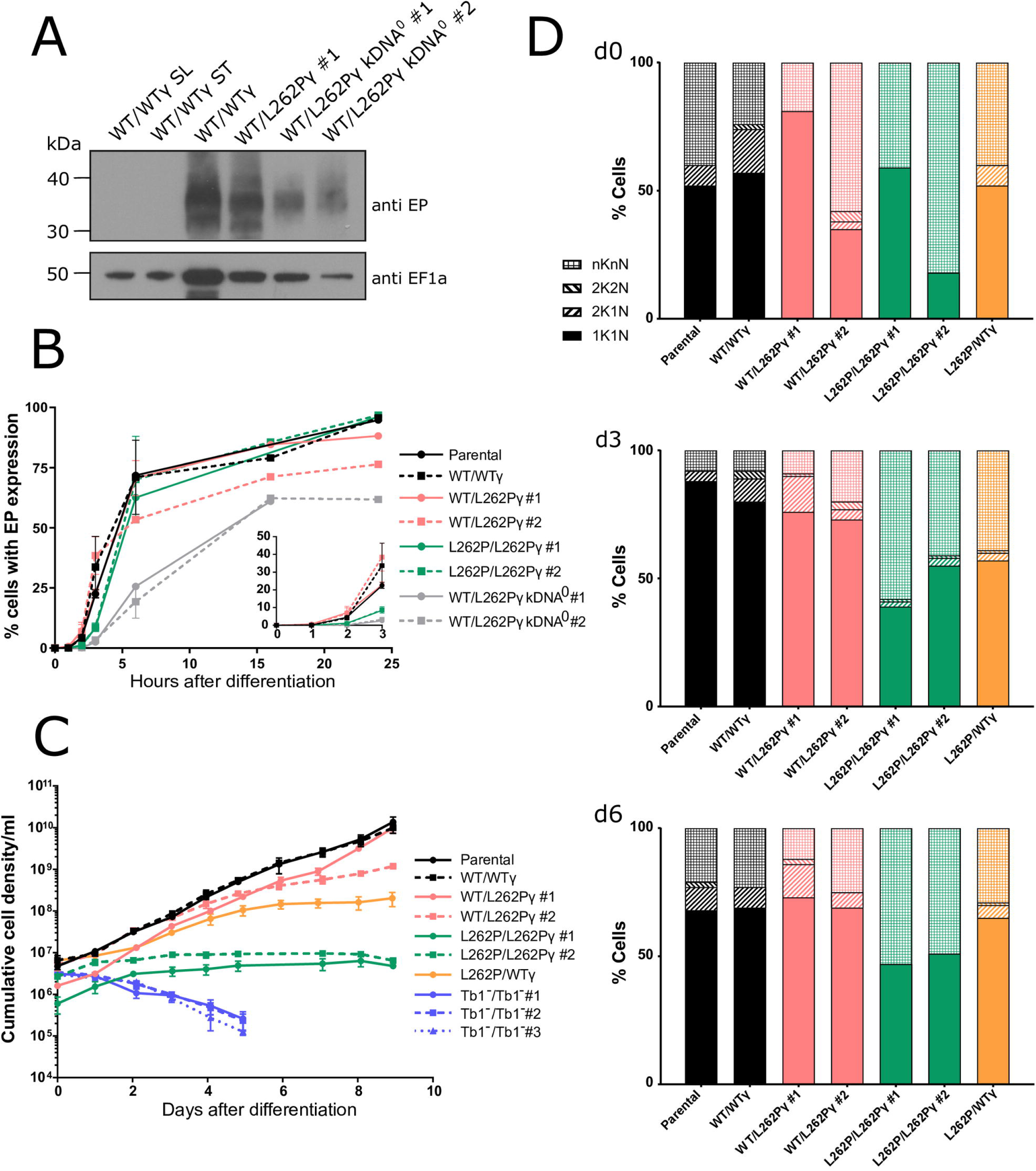
Ability of kDNA^0^ cells and F_1_F_O_-ATP synthase mutants to differentiate into PCF *in vitro*. (A) Western blot to assess EP procyclin expression. Stumpy cells were treated with 6 mM cis-aconitate (CA) and incubated at 27°C to induce differentiation. Cells were harvested after 24 h, 2×10^6^ cells were loaded per lane of an SDS-PAGE gel, and blots were probed with anti-EP procyclin (showing an expected smear due to heterogenous nature of EP protein, top panel) and anti-EF1α as a loading control (bottom panel). Slender and uninduced stumpy WT/WTγ cells were analysed as negative controls for EP expression. (B) EP expression measured by flow cytometry. Stumpy cells were treated with 6 mM CA and incubated at 27°C at the 0 h time point. At each time point, cells were fixed and stained with mouse anti-EP procyclin and anti-mouse IgG Alexa 488 antibodies. 10000 cells were analysed per sample; n=2. Inset plot magnifies the time points taken between 0-3 h after differentiation. Note that for kDNA^0^ cells, numbers at the 24 h time point represent large amounts of cell debris that stained positive for EP. No intact and motile cells were visible upon inspection by microscopy at this time point. (C) The growth of newly differentiated PCF *T. brucei*. Stumpy form cells were treated with 6 mM CA in HMI-9 medium for 24 h at 27°C. Cells were transferred to SDM80 medium at 27°C (t = 0 h) and cell density was determined with a haemocytometer over the course of 24 h. Counting was performed in triplicate with three separate blood cryostocks of each cell line thawed and differentiated at the same time. Error bars indicate standard deviation. kDNA^0^ cells were not included in this analysis due to their death within 24 h after induction of differentiation. (D) Cell cycle analysis of newly differentiated PCF *T. brucei*. Slides were prepared at days 0 (d0), 3 (d3) and 6 (d6) after differentiation and transfer into SDM80, and cells were blindly scored for cell cycle stage. Approximately 100 cells were staged per strain and time point. Category nKnN includes zoids (i.e., cells with no nucleus), monsters (multiple nuclei) and dying (large, blurry and/or fragmented nuclei) cells. kDNA^0^ and Tb1^-^ /Tb1^-^ cells were not included in this analysis due to their non-viability after differentiation.

We performed a time course to compare the kinetics of EP procyclin expression after induction of differentiation with CA in cells with and without kDNA. We collected samples from each cultured cell line at 0, 1, 2, 3, 6, 16 and 24 h after CA induction (+CA), stained them with mouse anti-EP procyclin antibody and anti-mouse IgG Alexa488, and analysed them by flow cytometry (Fig 1B). Parental, WT/WTγ and WT/L262Pγ cells containing kDNA expressed EP procyclin with similar kinetics; 55-70% of cells expressed EP procyclin by 6 h post induction. Two clones (#1 and #2) of the L262P/L262Pγ cell line initially showed slower kinetics of EP procyclin expression (Fig 1B inset), but, after 6 h, the percentage of EP procyclin positive cells was similar to other cell lines with kDNA. kDNA^0^ cells, on the other hand, showed a delayed rate of EP procyclin expression, with the proportion of cells expressing the protein increasing to a maximum of 55% by 16 h. As the vast majority of kDNA^0^ cells had died by 24 h after CA induction, this figure included EP procyclin-positive cell debris, which was not gated out of this analysis.

To test if freshly differentiated PCF cells expressing one or two of the mutated L262Pγ alleles could re-enter the cell cycle and proliferate under conditions that require efficient oxidative phosphorylation, we resuspended cells in SDM80 medium after 24 h of CA treatment. SDM80 lacks added glucose and contains N-acetyl D-glucosamine to inhibit uptake of residual glucose from FCS; it is in this low-glucose environment that PCF depend on oxidative phosphorylation to generate ATP [28, 37]. Here, ‘day 0’ (d0) was defined as being after 24 h in HMI-9 at 27°C plus CA, and then 24 h in SDM80 at 27°C. In these conditions, freshly differentiated WT/WTγ and WT/L262Pγ cells were able to proliferate (Fig 1C), and analysis of the number of kinetoplasts (K) and nuclei (N) per cell confirmed that they could progress through the cell cycle (Fig 1D). Cells with two kinetoplasts and one nucleus (2K1N) or with two kinetoplasts and two nuclei (2K2N) were consistently seen in culture across the time course, indicating actively dividing cells. Homozygous L262P/L262Pγ cell lines, however, did not grow in this medium. Although these cells survived in these conditions, they did not thrive and merely maintained the population density (observation by microscopy revealed that these cells were motile; data not shown). The L262P/L262Pγ cells were unable to progress through the cell cycle, with barely any 2K1N or 2K2N cell types detectable, and with noticeably less motility evident throughout the time course.

Cells of category nKnN, which included zoids (nK0N), ‘monsters’ with more than two nuclei and/or kinetoplasts, and other dying cells, were visible in this analysis at every time point and with every cell genotype (Fig 1D). This was possibly caused by cell swelling and/or nuclear fragmentation due to osmotic stress, which is a side effect of transferring freshly thawed stumpy cells from blood into liquid media for the differentiation experiments. Noticeably, for the L262P/L262Pγ cell lines, the proportion of nKnN cells remained extremely high as the time course continued, consistent with the high proportion of dying cells in these populations.

To confirm that the L262P/L262Pγ genotype, and not a secondary deficiency, was responsible for the severe growth defect of these PCF cell lines, we generated an L262P/WTγ (WTγ ‘add back’) cell line by replacing one subunit γ allele in cell line L262P/L262Pγ #2 with a WT subunit γ gene. The ability to maintain growth in SDM80 and progress through the cell cycle was partially restored in the resulting heterozygous γ cell line (Fig 1C, D), thus confirming an effect specifically due to the L262Pγ mutation.

We then asked whether 10 mM glucose in the medium (SDM79 medium) could rescue the growth defect of homozygous L262Pγ PCF cells, as ATP production via oxidative phosphorylation was reported to be inessential under these conditions [26,27,108], although this may be cell line dependent [98]. Surprisingly, we observed an immediate, severe growth phenotype in all freshly differentiated cell lines, with L262Pγ homozygotes having the strongest growth defect (S1A Fig). In WT/WTγ or heterozygous WT/L262Pγ cells, this phenotype could be rescued by removal of glucose from the medium, and restored by addition of glucose (S1B, C Fig). This growth retardation was probably due to an impact of glucose signalling on metabolic adaptation in freshly differentiated PCF parasites [109].

Cells lacking kDNA and homozygous L262Pγ mutants differ in two key aspects: due to the absence of all kDNA-encoded genes, the former cannot generate functional respiratory complexes III (cytochrome *bc_1_* complex) and IV (cytochrome *c* oxidase), nor the F_O_ part of the F_1_F_O_-ATP synthase. In the latter, the F_1_ and F_O_ parts are thought to be functionally uncoupled, although the extend of uncoupling is unclear (Table 1). To further investigate the basis for the phenotypic differences between these cell lines, we generated a cell line lacking ATP synthase subunit Tb1 (Tb1^-^/Tb1^-^; S2 Fig). Tb1 (systematic TriTrypDB ID Tb927.10.520) is the largest *T. brucei* F_O_ subunit and required to maintain the intact F_1_F_o_ complex structure and ATP synthase activity; however, it is not required for F_1_-ATPase activity [98, 99]. Like other F_O_ subunits, the gene becomes dispensable in BSF *T. brucei* in the presence of an L262Pγ allele, which enabled us to generate mutants that completely lack F_O_ but possess all kDNA-encoded genes (SFig 2). Three independent Tb1 null clones (Tb1^-^/Tb1^-^ #1, #2, #3) were used in this study. This cell line was able to generate stumpy forms *in vivo*, but once differentiated to the PCF *in vitro* they died rapidly (Fig 1C). This suggests that ATP production by oxidative phosphorylation becomes critical upon differentiation into insect stage parasites, and that the L262Pγ mutation only partially uncouples the F_1_F_O_-ATP synthase, permitting cell survival, if not proliferation (Fig 1C), via residual oxidative phosphorylation activity.

We next investigated the efficiency of mitochondrial ATP production pathways in the different cell lines. We added metabolic substrates that allowed ATP production by oxidative phosphorylation only (succinate) or a combination of oxidative and substrate phosphorylation (succinate plus pyruvate; α-ketoglutarate) to crude preparations of disrupted cells containing intact mitochondria. We then measured ‘*in organello*’ ATP production via the generation of a proportional luminescent signal [110]. Using succinate as a substrate, preparations from L262P/L262Pγ cells generated significantly less ATP when using succinate as a substrate than samples from cells with at least one WTγ allele (Fig 2A). Indeed, inhibition of the F_1_F_O_-ATP synthase with azide reduced ATP production in L262P/L262Pγ cell lysates only insignificantly, suggesting very minor, if any, contribution to ATP production by the mutated enzyme under these conditions. Preparations of WT/L262Pγ cells produced higher ATP levels than those of WTγ cells; at present we have no explanation for this observation. The presence of one or two L262Pγ alleles did not affect the level of representative F_1_ and F_O_ ATP synthase subunits (SFig 3), thus confirming the effect of the mutation is due to functional impairment. Adding back a WTγ allele to L262P/L262Pγ nearly restored ATP production via oxidative phosphorylation to levels found for WT/WTγ cells (Fig 2A). Interestingly, adding pyruvate as an additional substrate, which theoretically should allow additional ATP production via the ASCT cycle, did not significantly increase ATP production in mitochondria of any of the cell lines. This suggests the absence or low levels of either important transporters, enzymes or co-factors in these early differentiated cells. Similarly, α-ketoglutarate as sole substrate, which should allow ATP production by substrate phosphorylation via succinyl-CoA-synthetase, with production of succinate and subsequent ATP production via oxidative phosphorylation, resulted in only a moderate production of ATP in mitochondria from newly differentiated WTγ cells.

**Figure 2.**
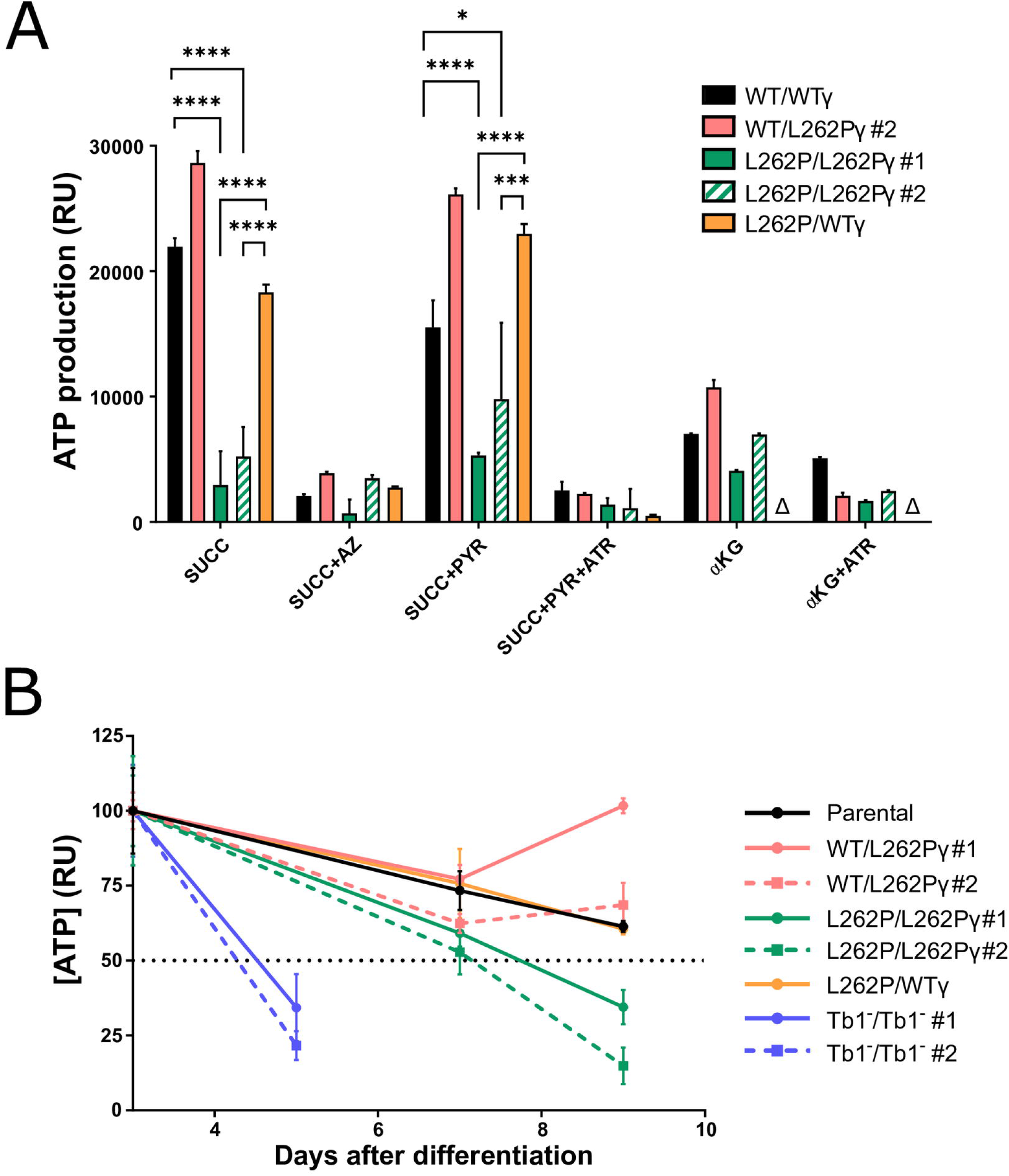
Mitochondrial ATP production capacity in F_1_F_O_-ATP synthase mutants. Cells were differentiated to PCF as described for Fig 1 and maintained in SDM80 medium. (A) Mitochondrial ATP production assay using permeabilised differentiated PCF cells with intact mitochondria. The capacity for ATP production via oxidative phosphorylation (succinate, SUCC), substrate level phosphorylation via the acetate:succinate CoA transferase (ASCT) / succinyl-CoA synthetase (SCoAS) cycle (pyruvate, PYR), and substrate level phosphorylation via α-ketoglutarate dehydrogenase and SCoAS (αKG) was assessed. Azide (AZ) and atractyloside (ATR) are inhibitors of the F_1_F_O_-ATP synthase and AAC, respectively. All assays were performed in triplicate at day 2 post differentiation. Error bars indicate standard deviation. Δ = assay not performed. P-values were determined using one-way ANOVA analysis. * < 0.05, ** < 0.01, *** < 0.001, **** < 0.0001. (B) Total cellular ATP level assays performed in triplicate, with the assay performed at days 3, 5, 7 and 9 post differentiation. Cell line Tb1^-^/Tb1^-^ was only assayed up to day 5 as insufficient numbers of viable cells remained after that time point. Error bars indicate standard deviation. 100% set to the ATP level from day 3 for each cell line, respectively.

The maintenance of total cellular ATP levels post differentiation was also impacted by the presence of homozygous L262Pγ. By day 9, the level of cellular ATP in L262P/L262Pγ cells was considerably lower than that of cells expressing at least one WTγ allele (Fig 2B). This is consistent with an impairment of ATP synthase complexes with an L262Pγ mutation in coupling the ΔΨm to ATP synthesis, as suggested above. ATP production assays could not be performed on Tb1^-^/Tb1^-^ cell lines due to the large number of viable differentiated cells required, but measurement of cellular ATP levels at day 3 showed that ATP levels were much depleted in these dying cells (Fig 2B).

Microscopic observation of L262P/L262Pγ cells grown in SDM80 medium had suggested a motility phenotype. We decided to investigate this further by recording motility tracks from videos of freshly differentiated cells in SDM80 medium. L262P/L262Pγ cells showed an evident motility defect in comparison with cells expressing at least one WTγ allele (Fig 3A). The average curvilinear velocities measured over the actual point-to-point route followed by the cell, i.e., the mean instant speeds, were calculated from these tracks, which confirmed that L262P/L262Pγ cells had a substantial progressive velocity defect manifesting between day 6 and day 9 post differentiation (Fig 3B). The WTγ add-back cell line L262P/WTγ, however, had a motility similar to that of WT/WTγ and WT/L262Pγ cells. Consistent with the more rapid drop in cellular ATP in Tb1^-^/Tb1^-^ cells, these mutants showed a pronounced reduction in mean velocity immediately after differentiation, at a time where the velocity of L262P/L262Pγ cells was still unaffected (Fig 3C, SFig 4). The rapid death of the Tb1^-^/Tb1^-^ cells after differentiation precluded an analysis of later time points.

**Figure 3.**
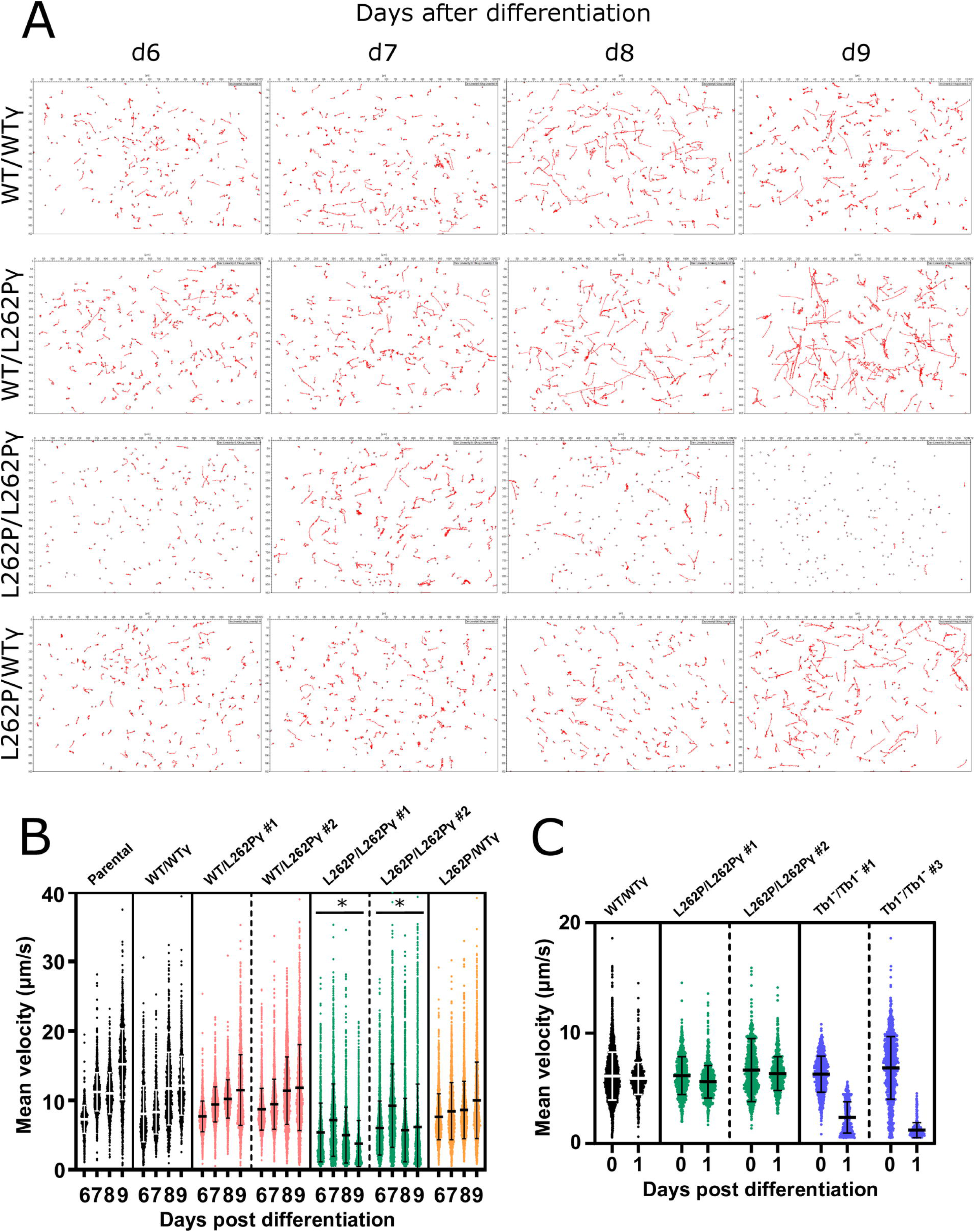
Motility analysis of *in vitro* cultured cells. (A) Representative cell movement tracks taken from videos of differentiated PCF cells *in vitro*. Videos were taken of populations of newly differentiated PCF *T. brucei* at days 6, 7, 8 and 9 (d6-9) post differentiation. (B) Measurement of curvilinear velocity (VCL) from video tracking in Fig 3A and expressed as ‘mean velocity’ of each tracked cell. VCL was assessed at days 6 - 9 (d6-9). At least 965 cells were tracked per cell line and time point. P-value determined using ANOVA analysis between cell lines across entire d6-d9 data set. Here, * indicates cell lines showing significant reduction in motility in comparison to the other cell lines shown (p < 0.05). Error bars indicate standard deviation. (C) VCL assessed as mean velocity of each tracked cell at days 0 (d0) and 1 (d1). At least 750 cells were tracked per cell line and time point. Error bars indicate standard deviation. Representative tracks taken from videos of these cells are shown in SFig 4.

To test whether trypanosomes devoid of kDNA could establish a PCF midgut infection *in vivo*, we fed teneral tsetse flies with horse blood containing stumpy form WT/L262Pγ kDNA^0^ parasites or kDNA^+^ stumpy form parasites with genotypes ‘parental’, WT/WTγ, WT/L262Pγ or L262P/L262Pγ. At day 9 post infection, we dissected the flies, and isolated, disrupted, and inspected their midguts under a microscope to assess the parasite infection prevalence. The parental cell line was able to differentiate to the PCF and established a midgut infection in around 85% of flies dissected (Fig 4A), with 70% of the flies having high levels of infection. In contrast, kDNA^0^ *T. brucei* cell lines were unable to establish even a single midgut infection.

**Figure 4.**
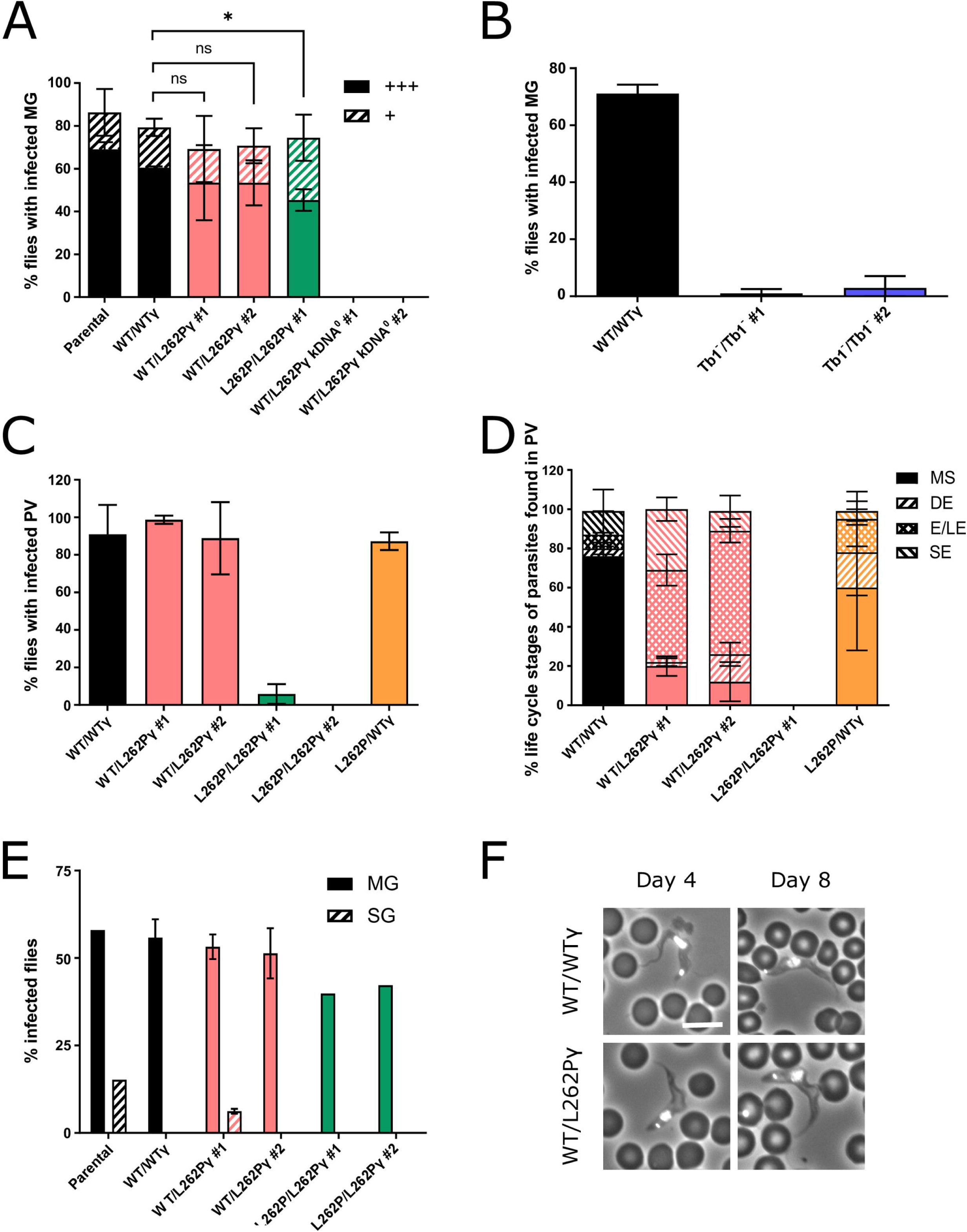
Assessment of the fly infectivity of cells with ATP synthase and kDNA mutations. (A) Infected tsetse fly midguts (MG) were harvested at day 9 post infected bloodmeal. Density of infection was judged by microscopy: +++ = % of flies found with a high level of midgut infection at >>10 parasites / field of view, + = a low level of midgut infection at ∼10 parasites / field of view. Approximately 25 flies were infected with one blood cryostock of stumpy form trypanosomes per cell line and replicate; n=3. Error bars indicate standard deviation. P-value determined by comparison between the heavy infection levels using unpaired t-test. * p < 0.05, ns = not significant (ns). Infections were performed at LSTM. (B) As in A, but approximately 50 flies were infected with one blood cryostock per cell line and replicate; n=3. Midguts (MG) were scored as infected if any trypanosomes were detected. Error bars indicate standard deviation. Infections were performed at LSTM. (C) Infected tsetse fly proventriculi (PV) were harvested between 21-28 days after infection. In total, 657 flies were dissected; n=3. Error bars indicate standard deviation. Infections were performed at Institut Pasteur. (D) Parasite life cycle stages of cells from infected PV in panel C were blindly quantified from 3 slides per cell line with approximately 100 cells per slide. L262P/L262P #2 was not assessed as no infected PV were detected (see panel C). MS = mesocyclic, DE = dividing epimastigote, E/LE = epimastigote/long epimastigote, SE = short epimastigote. Error bars indicate standard deviation. (E) Infected tsetse fly midguts (MG) and salivary glands (SG) were harvested four weeks after infection. For infections with WT/WTγ, WT/L262Pγ #1 and WT/L262Pγ #2 cell lines, n=2. For the remaining infections, n=1. Error bars indicate standard deviation. Around 200 flies were infected with one blood cryostock containing stumpy form trypanosomes per cell line and replicate. Infections were performed at LSTM. **(F)** Mice were infected with metacyclic form trypanosomes isolated from fly salivary glands from panel E. Tail snips were performed daily to assess parasitaemia, and DAPI-stained blood smears were collected on days stated. Scale bar indicates 10 μm.

Flies infected with WT/L262Pγ or L262P/L262Pγ cells had a slightly lower midgut infection level compared to flies infected with parental or WT/WTγ cells; approximately 70% were midgut-infected, but the difference in overall infection rates was not statistically significant (Fig 4A). However, flies infected with L262P/L262Pγ parasites showed a significantly lower proportion of highly infected midguts. Infections conducted at a second tsetse colony with a subset of the same cell lines showed more reduced levels of midgut infection for L262P/L262Pγ cells, and this phenotype was rescued in the WTγ add-back cell line (SFig 5). Flies infected in a separate experiment with Tb1^-^/Tb1^-^ cells produced infections barely detectable in the midgut (Fig 4B), which is consistent with their rapid death after differentiation *in vitro* (see Fig 2B). When combined, these results demonstrate that *T. brucei* requires an F_1_F_o_-ATP synthase complex capable of functioning in oxidative phosphorylation to produce an efficient tsetse midgut infection. A reduced capacity for oxidative phosphorylation, as demonstrated by the homozygous L262Pγ parasites, can sustain a midgut infection, but at a reduced rate. Absence of functional F_1_F_O_ ATP synthase, as in Tb1^-^/Tb1^-^ cells, permits minimal midgut infection, whereas absence of all kDNA-encoded products completely prevents midgut infection.

We then assessed whether the motility defect observed for L262P/L262Pγ parasites *in vitro* would affect the PV colonization of this mutant. Whereas a high proportion of midgut infections manifested into PV infections with cells having at least one WTγ allele, infections with L262P/L262Pγ cells produced either no (clone #1) or very low (clone #2) numbers of infected PV (Fig 4C). As predicted, adding back a WTγ allele to an L262P/L262Pγ cell line restored a normal PV infection rate. Infections performed at a second tsetse colony also produced substantially reduced PV infection rates for L262P/L262Pγ parasites that were also rescued in a WTγ add-back cell line (SFig 6A). Tracks recorded from cells released from the midgut showed that L262P/L262Pγ cells displayed significantly less motility than other cell lines within the midgut (Fig 5). We conclude that the low PV infection rate is correlated with a decreased motility of these cells.

**Figure 5.**
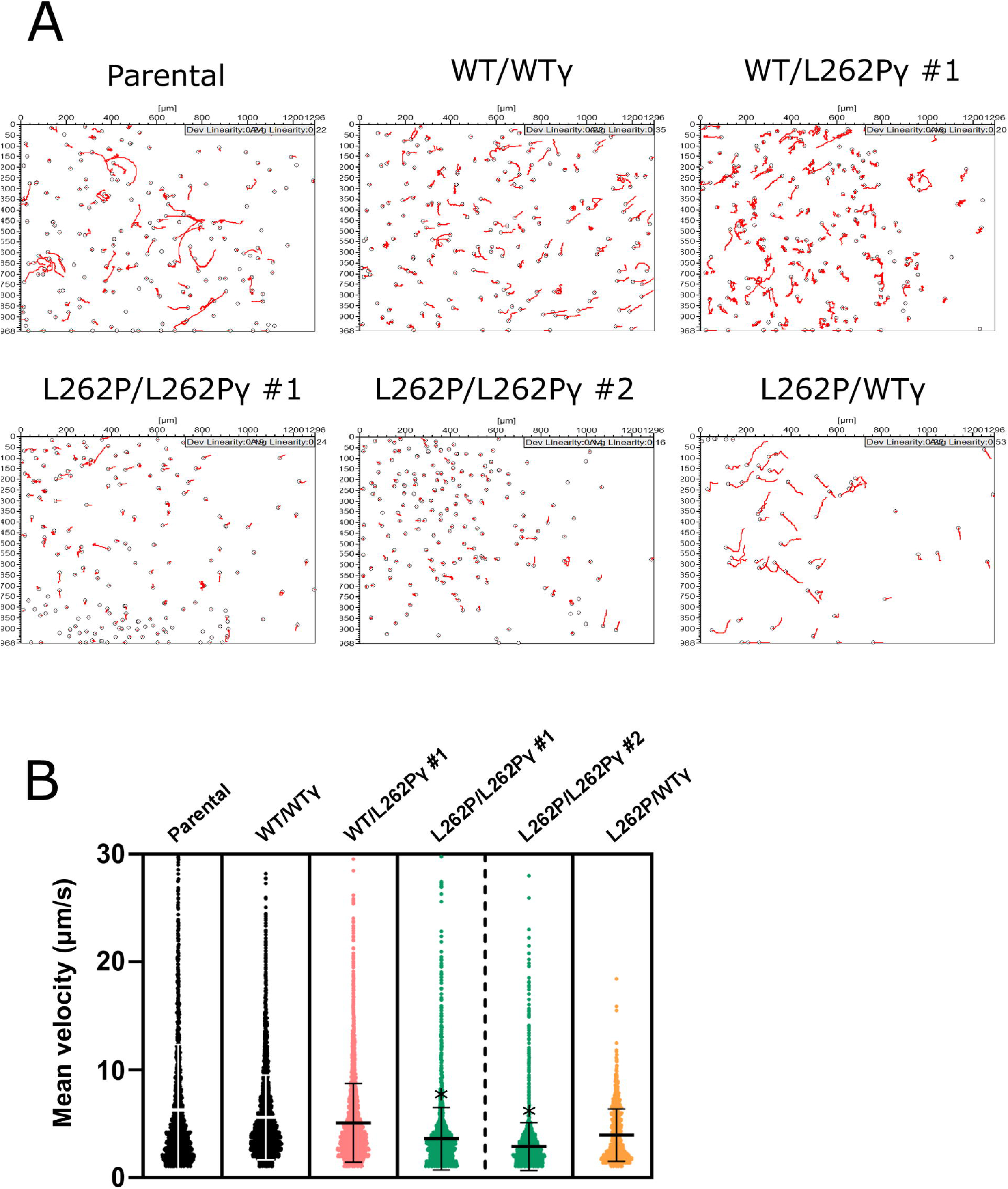
Motility analysis of parasites isolated from the fly midgut (*ex vivo*). (A) Representative tracks taken from videos of cells from tsetse fly midguts dissected at day 9 post infection at Institut Pasteur. (B) Measurement of curvilinear velocity (VCL, calculated as mean velocity for each cell) from tracked videos of *ex vivo* PCF cells. P-values were determined using ANOVA analysis between cell lines. Here, * indicates cell lines showing significant reduction in motility in comparison to the other cell lines; * p < 0.05. Error bars indicate standard deviation. At least 11 movies were taken per fly posterior midgut, with at least 3 flies dissected per strain. At least 1,546 cells were tracked per cell line.

To assess whether capacity for oxidative phosphorylation also affected differentiation, we quantified the various parasite stages found in the PV. All cells with at least one WTγ allele could differentiate into long mesocyclic trypomastigote forms as well as into long and short epimastigote forms (Fig 4D). WT/L262Pγ cells showed higher proportions of epimastigote stages than WT/WTγ cells, however introduction of a single WTγ allele in the L262P/L262Pγ background (L262P/WTγ) resulted in a percentage of epimastigote cells that was more similar to WT/WTγ. The life cycle stages of L262P/L262Pγ cells found in the PV were not assessed for experiments at this fly colony due to the extremely low level of infections found in this organ. The PV infection rate for the L262P/L262Pγ mutant at the second fly colony allowed us to assess the proportion of epimastigote forms for these cells and for parental and WTγ add-back cells as key controls. Interestingly, epimastigotes could be seen at approximately the same proportion in all these cell lines (SFig 6B), despite the severely reduced levels of PV infection with L262P/L262Pγ cells. This suggests that, if able to migrate to the PV, L262P/L262Pγ cells can physically progress to the epimastigote stage of the life cycle. Thus, whether a reduced capacity of these cells for oxidative phosphorylation affects the efficiency of differentiation to the epimastigote stage remains uncertain from these data.

To investigate whether the life cycle could be completed by L262Pγ-expressing cell lines, we dissected infected tsetse flies at 28 days post infection. Around 12% of tsetse fed with parental stumpy cells had detectable parasites in their salivary glands (Fig 4E), confirming the ability of this parental cell line to complete the *T. brucei* life cycle. WT/L262Pγ clone #1 produced a lower salivary gland infection rate at an average of 6%. No infected salivary glands were found in flies that had been fed WT/WTγ, WT/L262Pγ #2 or L262P/L262Pγ parasites, despite these flies having established midgut infections. The lack of salivary gland infections for the WT/WTγ and one of the WT/L262Pγ cell lines suggests that the capacity for this step in the life cycle progression is perhaps unstable and prone to loss in experimental cell lines. As a consequence, the significance of the lack of infectivity of L262P/L262Pγ cells is uncertain.

To confirm that the metacyclic form trypanosomes found within the salivary glands in the parental strain and in WT/L262Pγ #1 were indeed animal transmissible, we harvested these forms and used them to infect mice. Slender BSF trypanosomes were visible in both tail blood smears from day 4, with stumpy forms being visible in both strains from day 8 (Fig 4F). To ensure that cells of original genotype WT/L262Pγ had not reverted to WT/WTγ genotype, we confirmed by sequencing that both WT and L262P γ alleles were detectable at the DNA level (data not shown). Hence, *T. brucei* with a WT/L262Pγ genotype can complete their parasitic life cycle.

To the best of our knowledge, this is the first published example of a comparison of infection results from two distinct tsetse colonies. However, not all cell lines were challenged each time due to fly number restraints and to the extremely laborious nature of the experimental tasks. In total, result trends observed in parallel with the same strains were similar between the tsetse fly colonies at LSTM, Liverpool, and the Pasteur Institut, Paris. They only differed to the degree to which the L262P/L262Pγ mutation affected midgut and PV infections. A decrease in midgut infection level was seen for the Pasteur infections (SFig 5) when compared those performed at Liverpool (Fig 4A). This correlated with PV infection results, where the L262P/L262Pγ cells barely established a PV infection at Pasteur (Fig 4C), in contrast to Liverpool (SFig 6A), where more substantial PV infections were found, but at a lower level to those found in other cell types. These subtle differences could be explained by the different intrinsic vector competences of the two colonies and/or by differences in the infection and maintenance protocols (infection vehicle, bloodmeal type and frequency).

## Discussion

The kinetoplast (or kDNA) of the sleeping sickness parasite *T. brucei* encodes essential subunits of respiratory chain complexes, the F_1_F_O_ ATP synthase and the mitoribosome, which is a property it has in common with mitochondrial DNA of other eukaryotes. Although mitochondrial activity shows dramatic changes during the complex life cycle of the parasite, multiple *in vitro* studies have confirmed that the major proliferative stages in the mammalian bloodstream (long slender BSF) and in the tsetse fly vector midgut (PCF) depend on kDNA-encoded gene products to manufacture ΔΨm (BSF and PCF) and mitochondrial ATP (PCF). In the BSF, dependence on kDNA for viability can be overcome by compensatory changes in the nuclearly-encoded F_1_ subunit γ. We have utilised a panel of BSF mutants with various defects in kinetoplast-encoded functions (Table 1) combined with a number of multiple experimental challenges to address whether the compensatory mutation that functions in BSF can compensate for kDNA loss in PCF, and to what extent capacity for ATP production by oxidative phosphorylation is required for PCF differentiation and viability *in vivo*, as well as for cyclical development *in vivo* in the tsetse fly.

### Parasites heterozygous for the compensatory subunit γ mutation can complete the life cycle

A cell line heterozygous for the mutation that enables viability of kDNA^0^ BSF parasites, WT/L262Pγ #1, was able to complete the life cycle within the fly and produce animal-infective metacyclic parasites. Growth, capacity for oxidative phosphorylation and motility *in vitro*, and midgut infection prevalence all were unimpaired by the WT/L262Pγ genotype. Although WT/L262Pγ parasites displayed a higher rate of epimastigote form infection in the PV than WTγ parasites, this observation was probably not linked to the heterozygous genotype itself as another heterozygous cell line, the WTγ addback cell line (L262P/WTγ) showed levels of epimastigote forms that were more similar to the WTγ cell lines.

Approximately 25% of flies with parental strain midgut infections developed salivary gland infections (or ∼12.5% of total number of flies fed with parasites), which is consistent with infection levels with this strain reported elsewhere [111]. In flies infected with WT/L262Pγ #1 parasites, approximately 10% of flies with a midgut infection developed a salivary gland infection. However, as only the parental line and one WT/L262Pγ clone were able to differentiate into the metacyclic stage, and as the ability to invade the salivary glands seems to be a fragile trait that can be lost in laboratory adapted cell lines, it cannot be ruled out that these differences in infection rate could be due to clone-specific differences unrelated to the genetic manipulation. Nonetheless, our results clearly demonstrate that parasites with heterozygous L262Pγ mutations enabling kDNA independence can be transmitted by tsetse flies. *T. brucei* parasites with - this or other mutations that obviate the need for kDNA in BSF are much less sensitive to classes of drugs used in the field to treat animal disease, including isometamidium and homodium, since these drugs, at least in part, target parasite kDNA [112]. Exposure to drugs that target kDNA, however, would cause loss of kDNA from WT/L262Pγ cells, thus preventing transmission by tsetse flies as confirmed in the present study (see below). Hence, although a drug-resistant phenotype caused by a heterozygous subunit γ genotype could initially be spread by cyclical transmission, once in a host treated with these drugs, parasites would continue to proliferate, but the opportunity for further tsetse transmission from that host would stop. If such mutations were selected for in an animal-infectious strain of *T. b. brucei*, there would also be scope for sexually recombining with human-infective parasites to produce parasites with increased drug resistance able to infect humans [113, 114].

### The requirement for F_1_F_O_-ATP synthase complexes through the life cycle

A comparison of the phenotypes observed for cells with a homozygous L262P F_1_ subunit γ mutation vs. null mutants for F_O_ subunit Tb1 provided insight into the requirement for F_1_F_O_ complexes for life cycle completion. Firstly, L262P/L262Pγ cells (unlike WT/WTγ and WT/L262Pγ cells) grow poorly in glucose-depleted conditions that increase dependence on oxidative phosphorylation [28, 37]. Tb1^-^/Tb1^-^ cells, on the other hand, die rapidly under these conditions. Secondly, early in differentiation to PCF *in vitro*, L262P/L262Pγ cells maintain higher cellular ATP concentrations than Tb1^-^/Tb1^-^ cells. Thirdly, L262P/L262Pγ cells sustain midgut populations in the glucose-depleted tsetse fly midgut, although these are less dense and cells are less motile by day 9 than in flies infected with WT/WTγ or WT/L262Pγ cell lines. Tb1^-^/Tb1^-^ cells, in contrast, were unable to establish or sustain midgut infections. Together, these results demonstrate that, firstly, differentiation to PCF and tsetse infectivity depends on functional F_1_F_O_ ATP synthase complexes, which is consistent with the observation that stumpy parasites treated with the F_1_F_O_ inhibitor oligomycin were unable to re-enter the cell cycle during differentiation to PCF cells (Timms et al. 2002). Secondly, the L262Pγ mutation causes a less profound defect in F_1_F_O_-ATP synthase function than the loss of Tb1. This suggests that, whereas the F_1_F_O_-ATP synthase is nearly completely disrupted in the absence of Tb1 [98, 99], the F_1_F_o_ complex retains some functionality in L262P/L262Pγ cells as these cells can still generate enough ATP mitochondrially to survive in these conditions.

In principle, the effect of the mutations on mitochondrial ATP production could be direct (via F_1_F_O_-ATP synthase participation in oxidative phosphorylation), indirect (via an impact on substrate level phosphorylation) or a combination of both. Crude mitochondrial extracts from L262P/L262Pγ cells barely produce ATP via oxidative phosphorylation, with ATP production levels close to those in the presence of an ATP synthase inhibitor. This is consistent with the expected partial uncoupling between F_1_ and F_O_ parts that is caused by the disrupted interaction between the mutated γ subunit and F_1_ headpiece [84–87]. On the other hand, like stumpy cells [24,115,116], newly differentiated L262P/L262Pγ PCF cells are able to metabolise α-ketoglutarate via mitochondrial substrate level phosphorylation to generate ATP. Indeed, we found that L262P/L262Pγ #2 cells produce ATP from α-ketoglutarate to a similar extent as WTγ cells. As L262P/L262Pγ cells can establish infection in the tsetse midgut and progress to the PV, we postulate that the ability for substrate level phosphorylation in the absence of glucose may be higher *in vivo* than *in vitro*; threonine or glutamine concentrations, for example, may be higher per trypanosome in the midgut [117, 118]. The presence of symbiotic bacteria in the tsetse midgut and the influence of fly digestion may also impact the metabolism of midgut-associated parasites [35,119,120]. Nutrients may also be depleted faster within liquid culture than in the fly midgut.

Why would L262P/L262Pγ cells and Tb1^-^/Tb1^-^ cells differ in their capacity for mitochondrial ATP production via substrate level phosphorylation? In L262P/L262Pγ PCF cells, ΔΨm can be generated by electron transport chain components, namely respiratory complexes III and IV [44, 45]. In these cells, the partial uncoupling between the F_1_ and F_O_ parts would be expected to generate a proton leak through F_O_ into the matrix [84–87]. However, the viability of these PCF cells, and the fact that they can produce ATP in the presence of α-ketoglutarate, suggests that a ΔΨm of sufficient magnitude is maintained to support metabolite transport and mitochondrial protein import for substrate level phosphorylation. In Tb1^-^/Tb1^-^ cells, however, the near-complete loss of F_o_ from ATP synthase complexes [98, 99] might cause sustained hyperpolarisation of the inner mitochondrial membrane as the ΔΨm generated by complex III and IV cannot be dissipated by proton movement through the F_o_ moiety [121–123]. Hyperpolarisation would interfere with metabolite exchange and mitochondrial protein import and cause ROS generation and enhanced oxidative stress, thus damaging cells in a multitude of ways [124]. Indeed persistent hyperpolarisation is an acknowledged cell fate and death checkpoint [125–128]. In addition, the profound inability of Tb1^-^/Tb1^-^ cells to survive as PCF *in vitro* or *in vivo* may be influenced by the substantial remodelling of the mitochondrion required during BSF to PCF differentiation [9]. During this differentiation, the tubular BSF mitochondrion expands in volume and becomes elaborated into a branched network structure with abundant cristae [15,115,129], which are important for efficient oxidative phosphorylation [130–132]. Dimerization of the trypanosome F_1_F_o_ complex may be critical for folding of the inner mitochondrial membrane and cristae formation, and may critically involve F_O_, as in other organisms [133–138]. The absence of Tb1 and consequently, F_O_, in these cells could interfere with the remodelling of the inner mitochondrial membrane during differentiation, preventing the ordered processes required from being completed. The mutants with progressive developmental defects generated here therefore will be a useful resource for future studies aimed at correlating kDNA-encoded functions with the changes in mitochondrial physiology and ultrastructure that are a hallmark of trypanosome differentiation.

Although the reduced ATP production levels that we observed for L262P/L262Pγ cells are sufficient to permit colonisation of the midgut, they are correlated with a progressive motility defect that appears to decrease the migration efficiency of the cells in the fly. Only a low level of PV infection was detected in flies infected with L262P/L262Pγ cells. Swimming ability has been shown to be dictated by the availability of ATP for the dynein motor in numerous eukaryotic flagellate organisms [139–143]. In *T. brucei*, a dynein motor mutant has a motility defect in culture, and consequently is unable to migrate to the foregut and PV or infect the salivary glands [144]. Furthermore, efficient ATP generation is required for motility of *T. brucei* [145]. The ability to efficiently metabolise proline and alternative carbon sources may be particularly important as the parasite tries to migrate to compartments in the tsetse fly that could be less nutrient rich than the midgut, especially at the end of the digestive process [34, 35]. The lower infection level found in the L262P/L262Pγ infected midguts may also play a role in reduced migration as per the bottleneck model of *T. brucei* differentiation and migration in the tsetse fly [55].

In addition to motility, substantial amounts of ATP are also required for cell cycle progression and differentiation [146]. DNA replication complexes and kinesin proteins governing the vast morphological and organelle rearrangements depend on translocations fuelled by ATP [147–150]. Indeed, the progressive cellular ATP decline over time seen in L262P/L262Pγ cells is probably due to the large ATP expenditure necessary for motility and cell cycle progression, and the inefficiency of ATP generation in this cell line. *In vivo*, L262P/L262Pγ cells can differentiate into epimastigotes in a limited fashion, agreeing with recent evidence that oxidative phosphorylation may not be essential for this life cycle stage [35]. The presence of L262P/L262Pγ epimastigotes was unexpected considering that developmental progression in the insect may be driven by ROS production [63] and that the presence of a L262Pγ-induced proton leak would be hypothesised to neutralise ROS generation [151].

L262P/L262Pγ cell lines could not infect tsetse salivary glands despite these cell lines being able to sustain a midgut infection for over four weeks in these flies. The significance of this observation is uncertain, as it is common to observe a loss of fly infectivity in cultured *T. brucei* strains [152], as observed here for WT/WTγ and WT/L262Pγ #2 cell lines. As all cells had been maintained previously in BSF culture, it is possible that there was selection for parasites that had lost genes required for progression from the PCF in the tsetse midgut.

### The requirement for kDNA

Stumpy kDNA^0^ cells were consistently unable to (i) establish a midgut infection and (ii) differentiate into viable PCF cells *in vitro* after induction with CA, dying within 24 h. The kDNA^0^ stumpy cells express PAD1 [24], and do have a functional signalling pathway for differentiation as they respond to CA by expressing EP procyclin at a time point comparable to kDNA^+^ cell lines, as reported here and elsewhere [101, 104]. However, kDNA^0^ parasites do not effectively re-enter the cell cycle after differentiation from the cell cycle arrested stumpy BSF form. EP procyclin expression appears to be independent of kDNA, but effective re-entry into the cell cycle seems to either require the presence of the kinetoplast structure or a kDNA-encoded function, which is in agreement with Timms *et al.* (2002). We suggest that this is due to a complete failure to generate a ΔΨm. In the presence of functional kDNA, both the slender and stumpy BSFs generate ΔΨm by operating the F_1_F_o_-ATP synthase in the reverse direction [17–19,24,153]. The enzyme switches to the forward direction during stumpy to PCF differentiation [24, 154], and ΔΨm is then generated in a canonical fashion by respiratory complexes III and IV. According to the current model, kDNA^0^ long slender BSF can maintain ΔΨm independent of F_O_ via the electrogenic action of AAC-mediated ATP^4-^/ADP^3-^ exchange across the inner mitochondrial membrane when supported by efficient ATP hydrolysis via an F_1_ enzyme bearing a compensatory mutation in subunit γ [17,24,66,82,83]. However, ΔΨm is not generated in kDNA^0^ stumpy cells, probably due to mitochondrial substrate level ATP production using α-ketoglutarate preventing the electrogenic import of cytosolic ATP by the AAC [24]. In the absence of kDNA-encoded complex III and IV, kDNA^0^ PCF would be unable to generate ΔΨm for the same reason. We can conclude that all naturally occurring kDNA^-^ or kDNA^0^ variants of *T. brucei*, such as *T. b. evansi* and *T. b. equiperdum*, are intrinsically unable to undergo cyclical development in the tsetse.

## Materials and Methods

### Cell line generation

All cell lines used in this study were derived from culture-adapted pleomorphic *T*. *brucei* EATRO 1125 AnTat1.1 90:13 BSF parasites (S1 Table) [101]. Heterozygous cell lines with one F_1_F_O_-ATPase subunit γ allele (systematic TriTrypDB ID Tb927.10.180) with the L262P mutation (L262Pγ) and akinetoplastic (AK) versions of these cell lines were generated as detailed in [24]. Homozygous L262Pγ cell lines were generated by transfecting WT/L262Pγ *T. brucei* with plasmid pEnT6-γL262P-BSD (blasticidin resistance marker) to replace the remaining endogenous WTγ allele with an L262Pγ copy. The WTγ add back cell line was generated by releasing L262P/L262Pγ *T. brucei* cells from either puromycin or blasticidin selection before the transfection, and subsequently transfecting with plasmid pEnT6-γWT-PHL (phleomycin resistance marker). Both of these plasmids are based on the pEnT6 backbone [155] and contain either a L262Pγ gene or a wild type version (WTγ), facilitating the replacement of an ATPase γ subunit allele. The replaced gene is expressed by read-through transcription of the endogenous locus and contains its native 5’ untranslated region (UTR), but the aldolase 3’ UTR.

To generate a Tb1 (Tb927.10.520) null (Tb1^-^/Tb1^-^) cell line, pUC57-based plasmids containing either the blasticidin or the phleomycin resistance cassettes were designed and ordered from Biomatik (S2 Fig). Actin UTR 5’ and 3’ UTRs flank the resistance cassettes, with 350 bp of the Tb1 5’ UTR and intergenic region (IGR) positioned upstream of this, and 350 bp of the Tb1 3’ UTR and IGR positioned downstream of this. The first allele knock-out (KO) construct was designed to target sequences distal to those employed by the second allele KO construct [156]. These KO constructs were positioned inside *Hin*dIII restriction sites to enable digestion before transfection.

For the transfection, the AMAXA Nucleofector II (Lonza) was used with nucleofection solution (90 mM NaH_2_PO_4_, 5 mM KCl, 0.15 M CaCl_2_, 50 mM HEPES, pH 7.3) and program Z-001. *T*. *brucei* EATRO 1125 AnTat1.1 90:13 BSF clones were selected after 4 days at 37°C under drug selection as necessary (2.5 μg/ml G418, 5 μg/ml hygromycin, 0.1 μg/ml puromycin, 5 μg/ml blasticidin, 5 μg/ml phleomycin) in HMI-9 medium [157]; HMI-11 containing 10% (v/v) fetal calf serum (FCS; Gibco). ATPase γ genotypes were verified as detailed in Dewar et al., 2018. Tb1 KO cell lines were verified by PCR and western blot.

### *In vitro* differentiation and cell culture

Stumpy forms were generated in mice and were either used to make cryostocks containing a parasite population with approximately 90% stumpy bloodstream forms, or purified from blood [24], as required. *In vitro* differentiation was performed by adding 6 mM *cis*-aconitate (CA) to each culture, and cultures were left at 27°C for 24 h. If samples contained blood cells, the flasks were balanced on one corner to allow blood cells to settle out of the media. Newly differentiated PCF cells were washed and resuspended either in SDM79 containing 10 mM glucose (Invitrogen) [158] or in SDM80 at a density of at least 2×10^6^ cells per ml medium. SDM80 was made from SDM79 CGGGPPTA powder (PAA), an SDM79-based powder that lacks major carbon-containing components (sodium bicarbonate, glucose, glutamine, glutamate, proline, pyruvate, threonine and sodium acetate). All carbon sources except glucose were added back into the solution at the following concentrations: pyruvate 100 mg/l, L-proline 615 mg/l, L-threonine 394 mg/l, L-glutamine 320 mg/l, L-glutamate 24 mg/l, NaHCO_3_ 2 g/l, sodium acetate 10 mg/l. The media was supplemented with 50 mM N-acetyl D-glucosamine (Sigma) to prevent residual glucose uptake from FCS [159], and maintained under drug selection as necessary (15 μg/ml G418, 25 μg/ml hygromycin, 2 μg/ml puromycin, 10 μg/ml blasticidin, 2.5 μg/ml phleomycin). SDM79 and SDM80 were supplemented with 7.5 mg/l hemin and 10% FCS (Invitrogen), and cells were grown at 27°C continuously in these media, maintaining density over 2×10^6^/ml. For time courses, ‘day 0’ (d0) was defined as being after 24 h in HMI-9 at 27°C plus CA, and then 24 h in SDM80 at 27°C.

#### Protein expression analysis by flow cytometry and western blotting

For a quantitative measurement of EP expression, 2 x 10^6^ cells were harvested from +CA and –CA cultures at 0 h, 1 h, 2 h, 3 h, 6 h, 16 h and 24 h after CA addition. Upon collection, samples were transferred to 5 ml polystyrene round bottom tubes (BD Flacon 352052), centrifuged at 2000 x g for 5 minutes and washed in PBS (pH 7.4; 137 mM NaCl, 2.7 mM KCl, 10 mM Na_2_HPO_4_, 1.8 mM KH_2_PO_4_). The cell pellet was fixed in 500 µl PBS with 2% formaldehyde, 0.05% (w/v) glutaraldehyde overnight. Cells were pelleted, washed three times in PBS, and blocked in PBS with 2% (w/v) BSA for 1 h. After a PBS wash, cells were stained with PBS containing 2% BSA and 1/500 anti-EP procyclin monoclonal antibody (CedarLane Laboratories) for 1 h. Cells were washed in PBS, and then PBS containing 2% BSA and 1/1000 anti mouse IgG-Alexa 488 secondary antibody was added and left for 1 h. Cells were then washed and resuspended in 500 µl PBS containing 5 µg/ml Hoechst DNA staining dye (Life Tech.). After a 30 min incubation at room temperature, cells were analysed by flow cytometry at λ_ex_ 495 nm and λ_em_ 519 nm using a Beckton Dickinson LSRII machine with BD FACSDiva software and 2×10^4^ events per sample were measured. Results were analysed with FlowJo software (BD Biosciences).

Western blotting was performed as per Dewar et al., 2018. Antibodies used were anti-EP procyclin 1:500, anti-ATP synthase Tb1 1:2000 [160], anti-ATP synthase Tb2 1:2000 [160], anti-ATP synthase β 1:2000 [160], anti-COX IV 1:1000 [41], anti-COXVI 1:500 [44], anti-EF1α 1:7000 (Millipore), anti-mtHSP70 1:2000 [161].

### Tsetse fly handling and infections

To make the infective parasite stabilates, mouse blood containing stumpy form trypanosomes was harvested, mixed with 2% (w/v) sodium citrate to act as an anti-coagulant, and mixed 1:1 with HMI-9 containing 30% (v/v) FCS and 7.5% (v/v) glycerol for freezing at −80°C. Blood meals for tsetse fly infections were prepared as follows: one 200 µl aliquot per cell line was thawed at room temperature and mixed with ∼5 ml room temperature, sterile, defibrinated horse blood at the Liverpool School of Tropical Medicine (LSTM), or with ∼2 ml room temperature heat-inactivated foetal calf serum at the Institut Pasteur.

At the Liverpool School of Tropical Medicine, the *Glossina morsitans morsitans* (Westwood; origin Kenya) colony was maintained at 26±2°C and 68–78% relative humidity, with a 12 h light and 12 h dark cycle, and was fed triweekly on sterile, defibrinated horse blood in the dark using a sterile silicon membrane and a heated mat set to 37°C. All flies used in this study were newly emerged, teneral (unfed) male adults. Each experimental group was fed one infected bloodmeal containing predominantly stumpy form parasites when flies were 0–24 h after emergence. To obtain midgut infections, flies were offered a parasite-infected bloodmeal for 10 min, and then 24 h after feeding, flies were chilled to 4°C to remove unfed flies (identified by a non-scarlet abdomen). Flies with a visible bloodmeal were maintained at 27°C until dissection on day 9 post infection and fed uninfected bloodmeals as above every 2-3 days. For salivary gland infections, flies were allowed to feed on an infected bloodmeal for 15 min. Flies were chilled to 4°C at 24 h post bloodmeal to remove unfed flies. Flies were maintained as previously described and dissected four weeks post infection.

At the Institut Pasteur, *G. m. morsitans* tsetse flies (Westwood, origins Zimbabwe and Burkina Faso) were maintained in Roubaud cages at 27±1°C and 70±5% relative humidity, with a 12 h light and 12 h dark cycle, and fed twice a week through a silicone membrane with sterile, mechanically-defibrinated sheep blood at 37°C. Teneral males (between 24 h and 72 h post-emergence) were allowed to ingest BSF parasites in heat-inactivated fetal calf serum at 37°C through a silicone membrane for 10 min during their first meal and unfed flies were removed after feeding. When possible, flies were starved for at least 24 h before being dissected. For dissection, midguts and PVs were rapidly isolated and placed in distinct drops of phosphate buffer saline with 0.1% glucose as previously described [54, 61]. Isolated tissues were assessed by microscopy (40X) and imaged, or allowed to infuse for 5 min in a wet chamber before removal and collection of the remaining PBS drop containing trypanosomes for video recording (tracking analyses)[162].

To infect mice with metacyclic parasites harvested from tsetse flies, infected salivary glands were collected and pooled in ice-cold SDM79. Frozen stocks of these salivarian parasites were prepared by adding glycerol to a final concentration of 7.5% (v/v) before freezing at −80°C. Mice infections, blood smears and harvesting was performed as detailed in [24].

### Tracking and analysis (*in vitro* and *ex vivo*)

*In silico* 2D tracking was performed as previously described [144]. For each cell line, 10 to 20 movies were recorded for 20 seconds (50 ms of exposure). Trypanosomes freshly isolated from tsetse tissues and suspended in a drop of phosphate buffered saline with 0.1% (w/v) glucose and maintained at 27°C (*ex vivo*), or BSF trypanosomes sampled from cultures at 1×10^6^ parasites/ml and maintained at 37°C in matrix-dependent HMI-11 medium containing 0.5% methylcellulose (*in vitro*), were observed under the 10x objective of an inverted DMI-4000B microscope (Leica) coupled to an ORCA-03G (Hamamatsu) or a PRIM95B (Photometrics) camera. Movies were converted with the MPEG Streamclip V.1.9b3 software (Squared 5) and analysed with the medeaLAB CASA Tracking V.5.5 software (medea AV GmbH) for quantifying the mean velocity of 200 individual trypanosomes per movie over 149 successive frames. Statistical analyses and plots were performed with XLSTAT 2019.2.01 (Addinsoft) in Excel 2016 (Microsoft) or Prism V8.2.1 (GraphPad). Statistical analyses include two-sided ANOVA tests with Tukey ad-hoc post-tests for inter-group comparison at 95% confidence.

### *In organello* ATP production and whole-cell ATP quantification assays

The ATP quantification assay was performed as per the protocol for the CellTiter-Glo 3D cell viability assay kit (Promega), using 1x 10^4^ cells per sample. *In organello* ATP production assays were performed using the ATP Bioluminescence Assay kit CLSII (Roche) and digitonin extractions of 1 x 10^7^ cells per sample [110].

## Supporting information

**SFig 1. Effect of glucose on growth rates of freshly differentiated PCF cells.** (A) Cells were treated with 6 mM CA in HMI-9 for 24 h and then transferred to SDM80 medium supplemented with 10 mM glucose (solid lines) at a density of 2×10^6^/ml (day 0). An established PCF *T. brucei* cell line, 29:13 [163], was used as control that grows in either SDM80 (residual glucose, dashed lines) or SDM80 supplemented with glucose. Over the next 12-13 days, cells were counted once a day, and afterwards were diluted to a density of at least 3×10^6^/ml. (B, C) After induction of differentiation with CA, WT/WTγ (B) and WT/L262Pγ clone #2 parasites (C) were initially grown in either SDM80 (-glucose) or SDM80 supplemented with 10 mM glucose (+ glucose). Red arrows depict transfer of cells from SDM80 with glucose into SDM80 minus glucose (+/- glucose), and blue arrows depict transfer of cells into SDM80 minus glucose into SDM80 with glucose (-/+ glucose).

**SFig 2. Generation and verification of Tb1 null mutants.** (A) Tb1 knock-out plasmids produced by DNA synthesis (Biomatik). Drug selection markers (BSD, blasticidin resistance; PHL, phleomycin resistance) are flanked by actin 5’ and 3’ untranslated regions (UTR) for mRNA splicing and polyadenylation, respectively, and by Tb1 5’ and 3’ UTRs or intergenic regions (IGR) for homologous recombination. (B) PCR verification of three Tb1^-^/Tb1^-^ clones (#1, #2, #3). The top panel shows a duplex PCR with simultaneous amplification of a ∼700-bp fragment of the Tb1 coding sequence and a ∼900-bp fragment of the ATPase γ subunit gene as internal control. The bottom panel shows amplification of the Tb1 locus using primers flanking the 5’ and 3’ recombination sites. The wild type locus and the locus after replacement of the Tb1 coding sequence with the BSD and PHL genes give amplicons of 2400 bp, 2100 bp and 1540 bp, respectively. Genomic DNA from a WT/Tb1^-^ single knock-out cell line is included as a control. (C) Western blot verification of three Tb1^-^/Tb1^-^ clones, using a Tb1 antibody and an Hsp70 antibody as control. Whole cell lysates of 2×10^6^ cells were analysed per lane. The asterisk indicates non-specific detection of a ∼55-kDa protein by the Tb1 antibody. Lysates of an inducible Tb1 RNAi cell line (Hierro-Yap et al., 2021), uninduced and induced for 4 days for Tb1 ablation with tetracycline (tet), are shown as a control.

**SFig 3. Abundance of F_1_F_O_-ATP synthase in γ subunit mutants.** Cellular levels of mitochondrial respiratory complexes F_1_F_O_-ATP synthase and cytochrome oxidase (COX) assessed by probing a western blot with specific antibodies. Tb2 and β are subunits of the F_O_ and F_1_ moieties of the ATP synthase, respectively. COX is represented by subunits IV and VI. Cells were harvested after 48 h in SDM80 medium and whole cell lysates of 2×10^6^ cells were loaded per lane.

**SFig 4. Motility analysis of *in vitro* cultured cells.** Representative cell movement tracks taken from videos of differentiated PCF cells *in vitro*. Videos were taken of populations of newly differentiated PCF *T. brucei* at days 0 (d0) and 1 (d1) post differentiation, where d0 is defined as the timepoint after 24 h of exposure to 6 mM CA and 27°C and directly after transfer into SDM80 medium.

**SFig 5. Midgut infection rates at Institut Pasteur tsetse facility.** Infected tsetse fly midguts (MG) were harvested at day 9 after infection. Density of infection was judged by microscopy: +++ = % of flies found with a high level of midgut infection at >>10 parasites / field of view, + = a low level of midgut infection at ∼10 parasites / field of view. Infections were carried out with approximately 25 flies infected with one blood cryostock of stumpy form trypanosomes per cell line and replicate; n = 3. Error bars indicate standard deviation.

**SFig 6. PV infection rates at LSTM facility.** Dissections of tsetse fly PVs were conducted at four weeks post infection. Approximately 50 flies were infected with one cryovial of stumpy form trypanosomes per cell line. (A) The proportion of flies with infected midguts that had infected PV. Each set of infections was performed twice. (B) The proportion of trypanosomes with epimastigote morphology in the PV. Three infected PV per cell line were disrupted and left to air-dry on a slide. Slides were fixed and DAPI stained, and approximately 100 cells were staged per slide.

**S1 Table. Cells lines used in this study.**

## Notes

### Competing Interest Statement

The authors have declared no competing interest.

