## Supporting information for "Oxidative phosphorylation is required for powering motility and development of the sleeping sickness parasite *Trypanosoma brucei* within the tsetse fly vector"

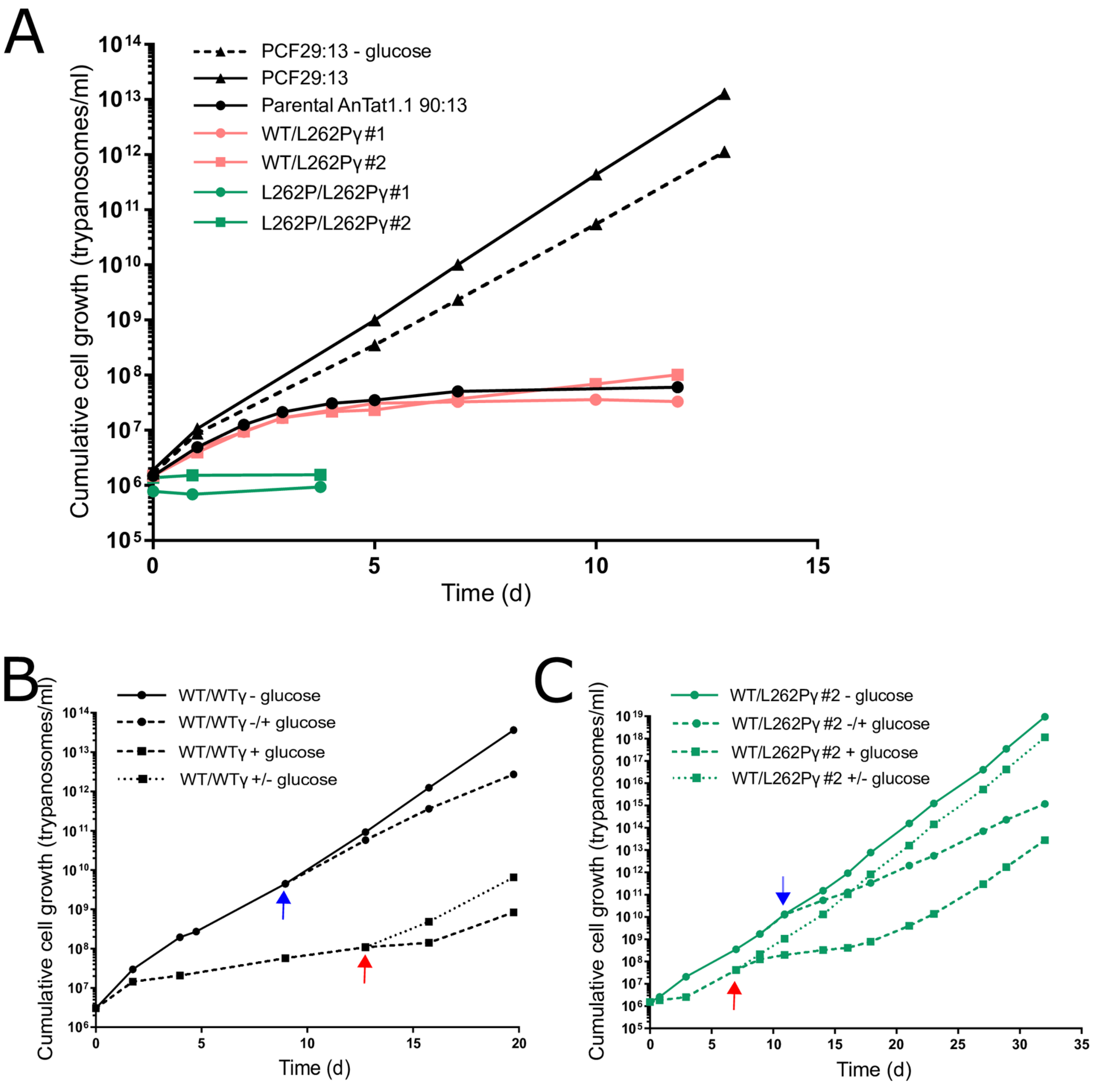


**SFig 1. Effect of glucose on growth rates of freshly differentiated PCF cells.** (A) Cells were treated with 6 mM CA in HMI-9 for 24 h and then transferred to SDM80 medium supplemented with 10 mM glucose (solid lines) at a density of $2x10^6$/ml (day 0). An established PCF T. brucei cell line, 29:13 [163], was used as control that grows in either SDM80 (residual glucose, dashed lines) or SDM80 supplemented with glucose. Over the next 12-13 days, cells were counted once a day, and afterwards were diluted to a density of at least $3x10^6$/ml. (B, C) After induction of differentiation with CA, WT/WTγ (B) and WT/L262Pγ clone #2 parasites (C) were initially grown in either SDM80 (- glucose) or SDM80 supplemented with 10 mM glucose (+ glucose). Red arrows depict transfer of cells from SDM80 with glucose into SDM80 minus glucose (+/- glucose), and blue arrows depict transfer of cells into SDM80 minus glucose into SDM80 with glucose (-/+ glucose).

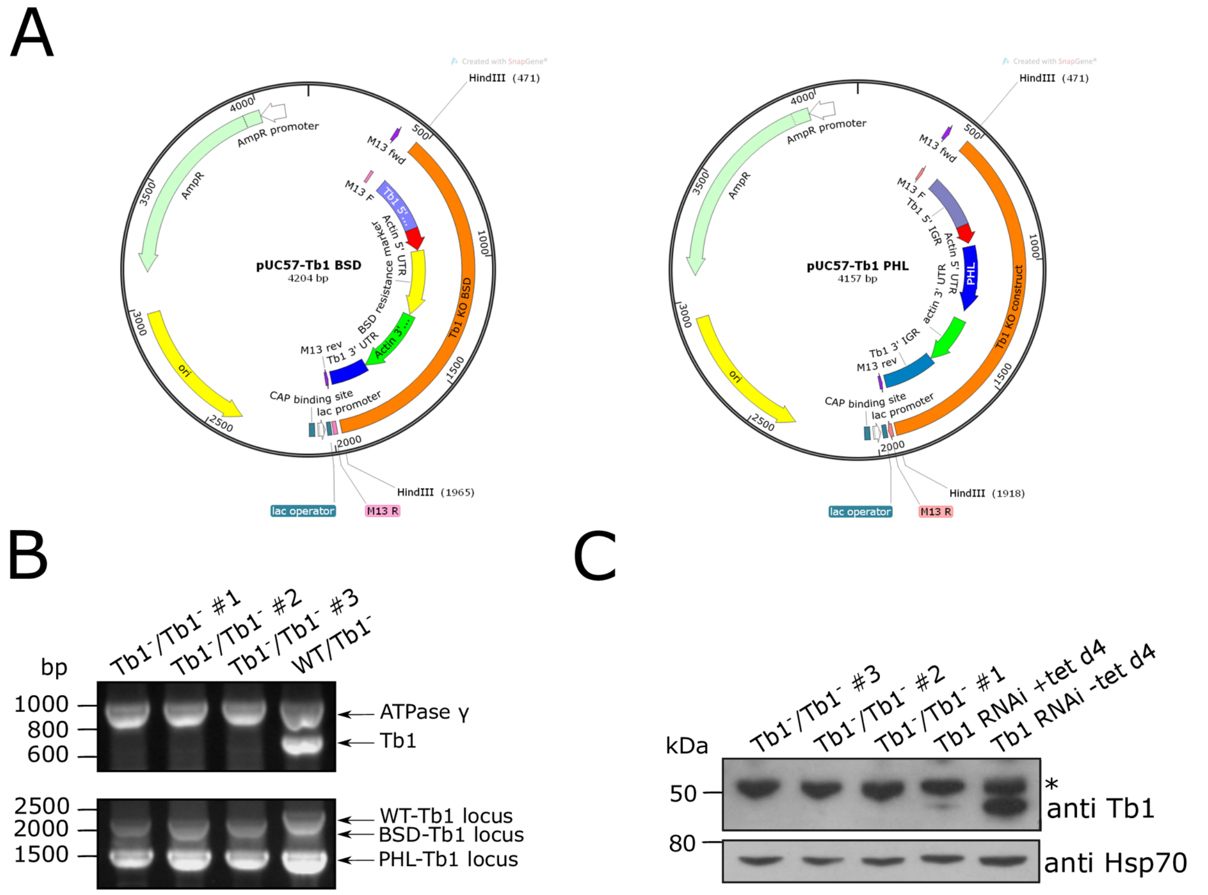


**SFig 2. Generation and verification of Tb1 null mutants.** (A) Tb1 knock-out plasmids produced by DNA synthesis (Biomatik). Drug selection markers (BSD, blasticidin resistance; PHL, phleomycin resistance) are flanked by actin 5' and 3' untranslated regions (UTR) for mRNA splicing and polyadenylation, respectively, and by Tb1 5' and 3' UTRs or intergenic regions (IGR) for homologous recombination. (B) PCR verification of three Tb1-/Tb1- clones (#1, #2, #3). The top panel shows a duplex PCR with simultaneous amplification of a ~700-bp fragment of the Tb1 coding sequence and a ~900-bp fragment of the ATPase γ subunit gene as internal control. The bottom panel shows amplification of the Tb1 locus using primers flanking the 5' and 3' recombination sites. The wild type locus and the locus after replacement of the Tb1 coding sequence with the BSD and PHL genes give amplicons of 2400 bp, 2100 bp and 1540 bp, respectively. Genomic DNA from a WT/Tb1- single knock-out cell line is included as a control. (C) Western blot verification of three Tb1-/Tb1- clones, using a Tb1 antibody and an Hsp70 antibody as control. Whole cell lysates of $2x10^6$ cells were analysed per lane. The asterisk indicates non-specific detection of a ~55-kDa protein by the Tb1 antibody. Lysates of an inducible Tb1 RNAi cell line (Hierro-Yap et al., 2021), uninduced and induced for 4 days for Tb1 ablation with tetracycline (tet), are shown as a control.

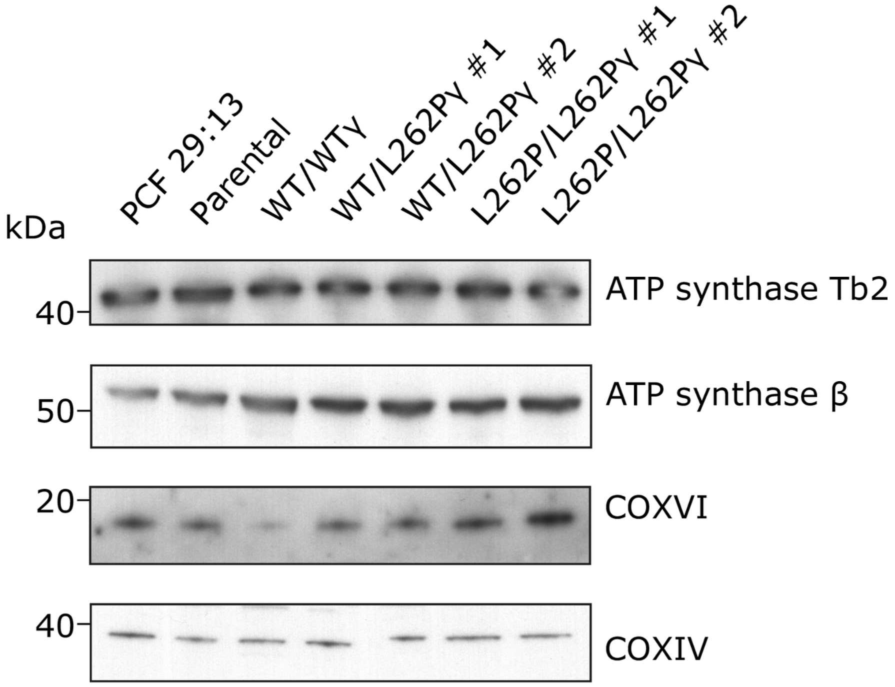


**SFig 3. Abundance of $F_1F_O$-ATP synthase in γ subunit mutants.** Cellular levels of mitochondrial respiratory complexes $F_1F_O$-ATP synthase and cytochrome oxidase (COX) assessed by probing a western blot with specific antibodies. Tb2 and β are subunits of the $F_O$ and $F_1$ moieties of the ATP synthase, respectively. COX is represented by subunits IV and VI. Cells were harvested after 48 h in SDM80 medium and whole cell lysates of $2x10^6$ cells were loaded per lane.

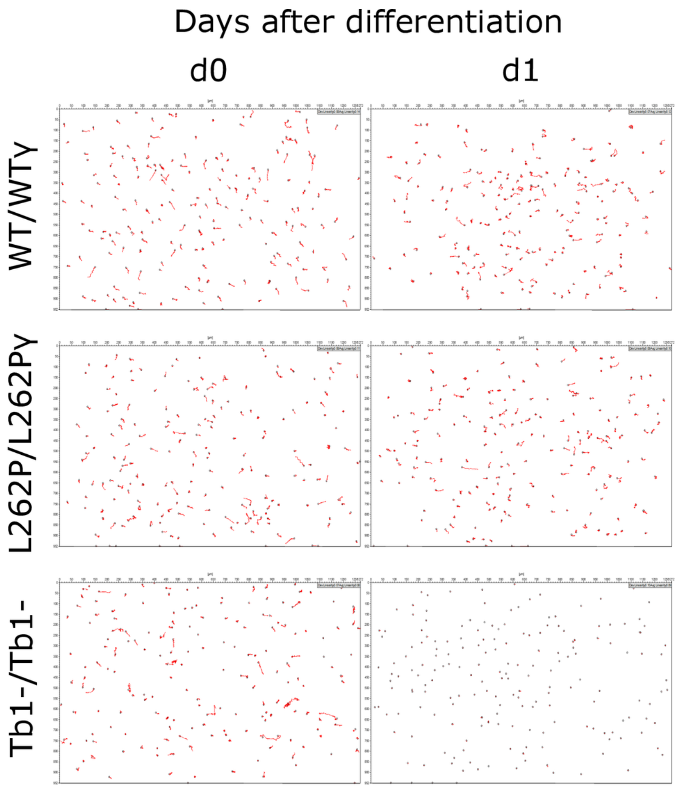


**SFig 4. Motility analysis of in vitro cultured cells.** Representative cell movement tracks taken from videos of differentiated PCF cells in vitro. Videos were taken of populations of newly differentiated PCF T. brucei at days 0 (d0) and 1 (d1) post differentiation, where d0 is defined as the timepoint after 24 h of exposure to 6 mM CA and 27°C and directly after transfer into SDM80 medium.

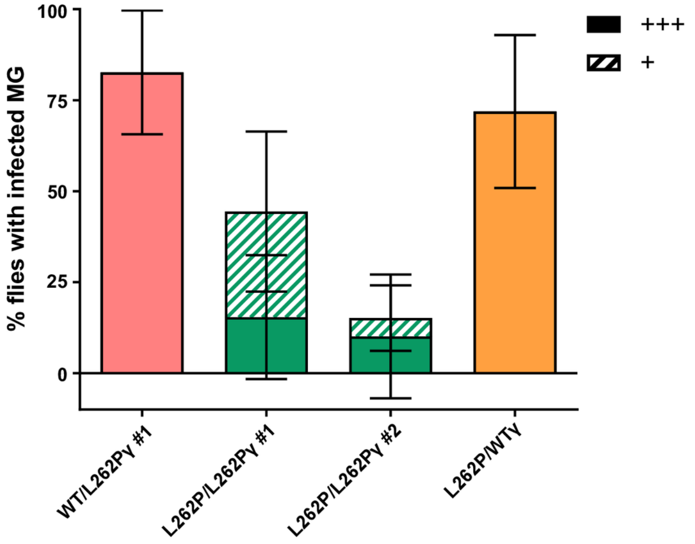


**SFig 5. Midgut infection rates at Institut Pasteur tsetse facility.** Infected tsetse fly midguts (MG) were harvested at day 9 after infection. Density of infection was judged by microscopy: +++ = % of flies found with a high level of midgut infection at >>10 parasites / field of view, + = a low level of midgut infection at ~10 parasites / field of view. Infections were carried out with approximately 25 flies infected with one blood cryostock of stumpy form trypanosomes per cell line and replicate; n = 3. Error bars indicate standard deviation.

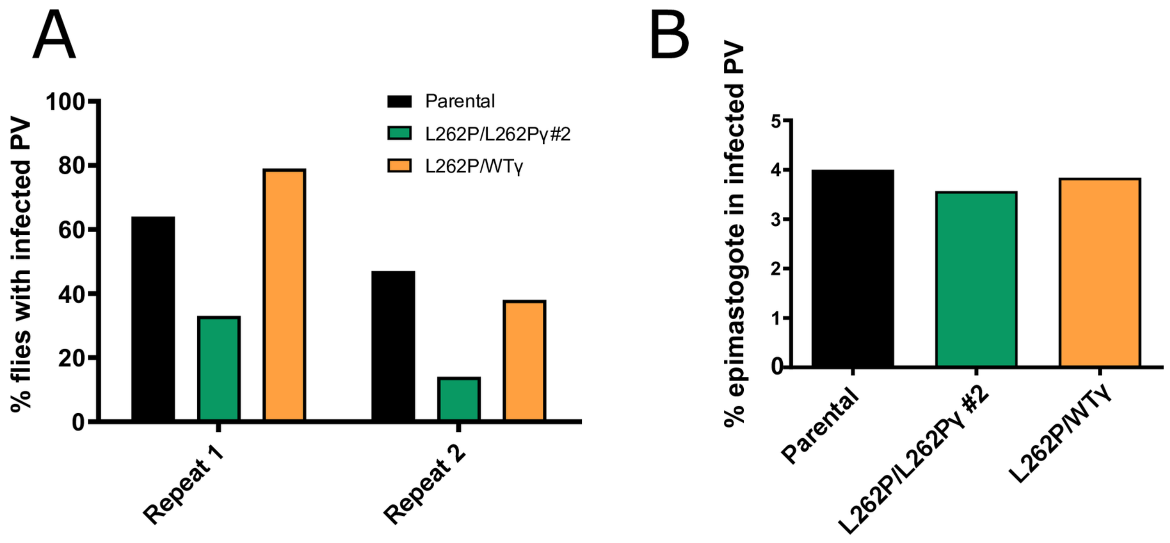


**SFig 6. PV infection rates at LSTM facility.** Dissections of tsetse fly PVs were conducted at four weeks post infection. Approximately 50 flies were infected with one cryovial of stumpy form trypanosomes per cell line. (A) The proportion of flies with infected midguts that had infected PV. Each set of infections was performed twice. (B) The proportion of trypanosomes with epimastigote morphology in the PV. Three infected PV per cell line were disrupted and left to air-dry on a slide. Slides were fixed and DAPI stained, and approximately 100 cells were staged per slide.

| Cell line | Derived from: | Genotype | Description |
|---|---|---|---|
| Parental | - | - | EATRO 1125AnTat 1.1 90:13 |
| WT/WTγ | EATRO 1125AnTat 1.1 90:13 | ATPγ/Δatpγ::atpγWT +PURO | One F1FO-ATP synthase subunit γ allele replaced with a WT copy and the puromycin resistance gene (PURO) |
| WT/L262Pγ #1 | EATRO 1125AnTat 1.1 90:13 | ATPγ/Δatpγ::atpγL262P +PURO | One F1FO-ATP synthase subunit γ allele replaced with a copy with the L262P mutation and the puromycin resistance gene (PURO) |
| WT/L262Pγ #2 | EATRO 1125AnTat 1.1 90:13 | ATPγ/Δatpγ::atpγL262P +PURO | One F1FO-ATP synthase subunit γ allele replaced with a copy with the L262P mutation and the puromycin resistance gene (PURO) |
| L262P/L262Pγ #1 | WT/L262P #3 | Δatpγ::atpγL262P +BSD/Δatpγ::atpγL262P +PURO | One F1FO-ATP synthase subunit γ allele replaced with a copy with the L262P mutation and the puromycin resistance gene (BSD) |
| L262P/L262Pγ #2 | WT/L262P #3 | Δatpγ::atpγL262P +BSD/Δatpγ::atpγL262P +PURO | One F1FO-ATP synthase subunit γ allele replaced with a copy with the L262P mutation and the puromycin resistance gene (BSD) |
| L262P/WTγ | L262P/L262P #2 | Δatpγ::atpγL262P +BSD/Δatpγl262p::atpγWT +PHL | One F1FO-ATP synthase subunit L262Pγ allele replaced with a WT copy with the puromycin resistance gene (PHL) |
| WT/L262PγkDNA0 #1 | WT/L262P #1 | ATPγ/Δatpγ::atpγL262P +PURO | Cell line WT/L262Pγ #1 induced to lose kDNA through exposure to acriflavine |
| WT/L262PγkDNA0 #2 | WT/L262P #2 | ATPγ/Δatpγ::atpγL262P +PURO | Cell line WT/L262Pγ #2 induced to lose kDNA through exposure to acriflavine |
| Tb1-/Tb1- #1 | WT/L262P #1 | ATPγ/Δatpγ::atpγL262P +PURO Δtb1::BSD/Δtb1::PHL | Cell line WT/L262Pγ #1 with BSD and PHL resistance genes present in Tb1 loci |
| Tb1-/Tb1- #2 | WT/L262P #1 | ATPγ/Δatpγ::atpγL262P +PURO Δtb1::BSD/Δtb1::PHL | Cell line WT/L262Pγ #1 with BSD and PHL resistance genes present in Tb1 loci |
| Tb1-/Tb1- #3 | WT/L262P #1 | ATPγ/Δatpγ::atpγL262P +PURO Δtb1::BSD/Δtb1::PHL | Cell line WT/L262Pγ #1 with BSD and PHL resistance genes present in Tb1 loci |

**STable 1. Cell lines used in this study.**
